# High throughput small molecule screening reveals NRF2-dependent and - independent pathways of cellular stress resistance

**DOI:** 10.1101/778548

**Authors:** David B. Lombard, William Kohler, Angela H. Guo, Christi Gendron, Melissa Han, Weiqiao Ding, Yang Lyu, Xinna Li, Xiaofang Shi, Zaneta Nikolovska-Coleska, Yuzhu Duan, Thomas Girke, Ao-Lin Hsu, Scott D. Pletcher, Richard A. Miller

**Affiliations:** Department of Pathology, University of Michigan, Ann Arbor, MI; Institute of Gerontology, University of Michigan, Ann Arbor, MI; Department of Molecular and Integrative Physiology, University of Michigan, Ann Arbor, MI; Department of Internal Medicine, Division of Geriatric and Palliative Medicine, University of Michigan, Ann Arbor, MI; Research Center for Healthy Aging, China Medical University, Taichung, Taiwan; Institute for Integrative Genome Biology, University of California Riverside, Riverside, CA

## Abstract

Biological aging is the dominant risk factor for most chronic diseases. Development of anti-aging interventions offers the promise of preventing many such illnesses simultaneously. Cellular stress resistance is an evolutionarily conserved feature of longevity. Here, we identify compounds that induced resistance to the superoxide generator paraquat (PQ), the heavy metal cadmium (Cd), and the DNA alkylator methyl methanesulfonate (MMS). Some rescue compounds conferred resistance to a single stressor, while others provoked multiplex resistance. Induction of stress resistance in fibroblasts was predictive of longevity extension in a published large-scale longevity screen in *C. elegans*. Transcriptomic analysis implicated Nrf2 signaling in stress resistance provided by two protective compounds, cardamonin and AEG 3482. Molecules that conferred stress resistance also induced cellular inflammatory pathways, and other core pathways such as AMPK signaling. Small molecules identified in this work may represent attractive candidates to evaluate for their potential pro-health and pro-longevity effects in mammals.

## Introduction

Aging is the key risk factor for most chronic debilitating diseases, conditions that impose great human suffering and ever-increasing health care costs in industrialized societies^1^. Genetic studies suggest that some of the pathways that govern the rate of aging and the onset of age-related disease in model organisms – in particular, insulin/IGF-like and mTOR signaling – may function similarly in humans to limit healthy lifespan^2–4^. A large body of evidence has demonstrated empirically that the aging rate can be dramatically slowed in invertebrates and in rodents^5^. Mice subjected to dietary restriction, or with certain single gene mutations, live much longer than controls^5^. Studies by the Interventions Testing Program (ITP) and other groups have shown that specific small molecules, such as rapamycin, acarbose, and 17α-estradiol, can substantially increase mouse lifespan^6–13^. In these studies, experimental animals typically remain healthy even late in life, with a much-reduced disease burden compared to controls. Thus, drugs with anti-aging activity appear to delay or abrogate many age-associated pathologies^14^.

To date, all compounds that delay aging in mammals have been identified by testing based on prior knowledge of mechanism of action, and/or data generated in invertebrate models, rather than via unbiased screening approaches. Unfortunately, the major vertebrate model used in aging biology, *Mus musculus*, has a lifespan of nearly three years, presenting a prohibitive barrier to large-scale direct screening for anti-aging effects using lifespan as an endpoint. The most extensive effort to identify compounds with anti-aging activity in mice, the NIA Interventions Testing Program, cannot test more than a handful of agents each year. Thus, although rodent studies have conclusively proven that extension of mammalian lifespan by small molecules is possible, currently only a handful of drugs with replicable lifespan benefits, *i.e.* > 10% extension, have been identified.

To circumvent this challenge, several groups have performed screens in more tractable, short-lived invertebrate model organisms to identify small molecules that increase lifespan. The first such large-scale screen, performed in the nematode roundworm *C. elegans,* revealed that certain serotonin signaling inhibitors extend longevity in this organism^15^. Other screens have revealed the ability of anticonvulsants^16^, angiotensin converting enzyme antagonists^17^, and modulators of other signaling pathways to increase worm lifespan^18, 19^. Recent studies have employed *in silico* strategies to prioritize compounds for direct lifespan testing in nematodes and other organisms^20–22^.

An alternative, and complementary, strategy to these approaches is to evaluate test compounds in mouse or human cells, using phenotypes that serve as surrogates for organismal lifespan. In this regard, cellular resistance to environmental stressors is a frequent correlate of longevity^23^. In mammals, dermal fibroblasts derived from longer-lived species, or from long-lived mouse mutants, show resistance to some forms of lethal injury^24–30^. Similarly, in *C. elegans,* many long-lived mutants show resistance to a variety of environmental insults^31^, including oxidative stress, a phenotype that has been used to screen for longevity mutants^32^. An analogous strategy, based on screens for surrogate stress resistance endpoints, has been employed to identify long-lived mutants in budding yeast^33^. A group of compounds that promote lifespan extension in worms was enriched for molecules that protect against oxidative stress^18^.

We performed high-throughput screening (HTS) to identify small molecules that induce resistance against multiple forms of cellular injury in fibroblasts derived from adult skin biopsies of genetically heterogeneous mice. Three screens were performed in parallel, based on resistance to the superoxide generator paraquat (PQ), the heavy metal cadmium (Cd), and the DNA alkylating agent methyl methanesulfonate (MMS). Since many of our hit compounds are already FDA-approved drugs with favorable safety profiles, the agents that induce stress resistance in our screens may warrant further testing in mice for health and longevity benefits, and subsequently could potentially be translated into clinical trials for anti-aging effects humans.

## Results

### HTS identifies small molecules that induce resistance against multiple stressors in mammalian fibroblasts

We performed HTS using mouse tail fibroblasts (MTFs). Stress resistance of this cell type correlates with organismal longevity in mouse strains in which single gene mutations extend healthy lifespan^24–29, 34–38^. We initially defined the basic parameters for cell-based stress assays using this cell type, optimizing growth media, seeding density, and incubation time. Using these parameters, we then performed an extensive series of pilot experiments using the cellular stressors PQ, MMS, and Cd, along with selected candidate chemical rescue agents (curcumin and arsenite), to optimize conditions for HTS for induction of stress resistance (Fig. S1A-C and data not shown).

Using these optimized conditions, we evaluated small molecule libraries containing a total of 6,351 compounds (Biofocus NCC, MicroSource Spectrum 2400, Prestwick, LOPAC, and a focused collection), including some duplicates included in more than one library, for ability to protect MTFs from cell death induced by PQ at a single dose, 16 μM (Table S1). In total, >4,500 unique compounds were screened. A typical assay is shown, in which 640 candidate rescue agents were tested (Fig. 1A). Many compounds within this particular set protected cells against PQ toxicity. Testing of duplicate plates of rescue compounds revealed low plate-to-plate variation (Fig. S1D). Similar primary screens were conducted using Cd and MMS as stressors (Table S1).

**Figure 1:**
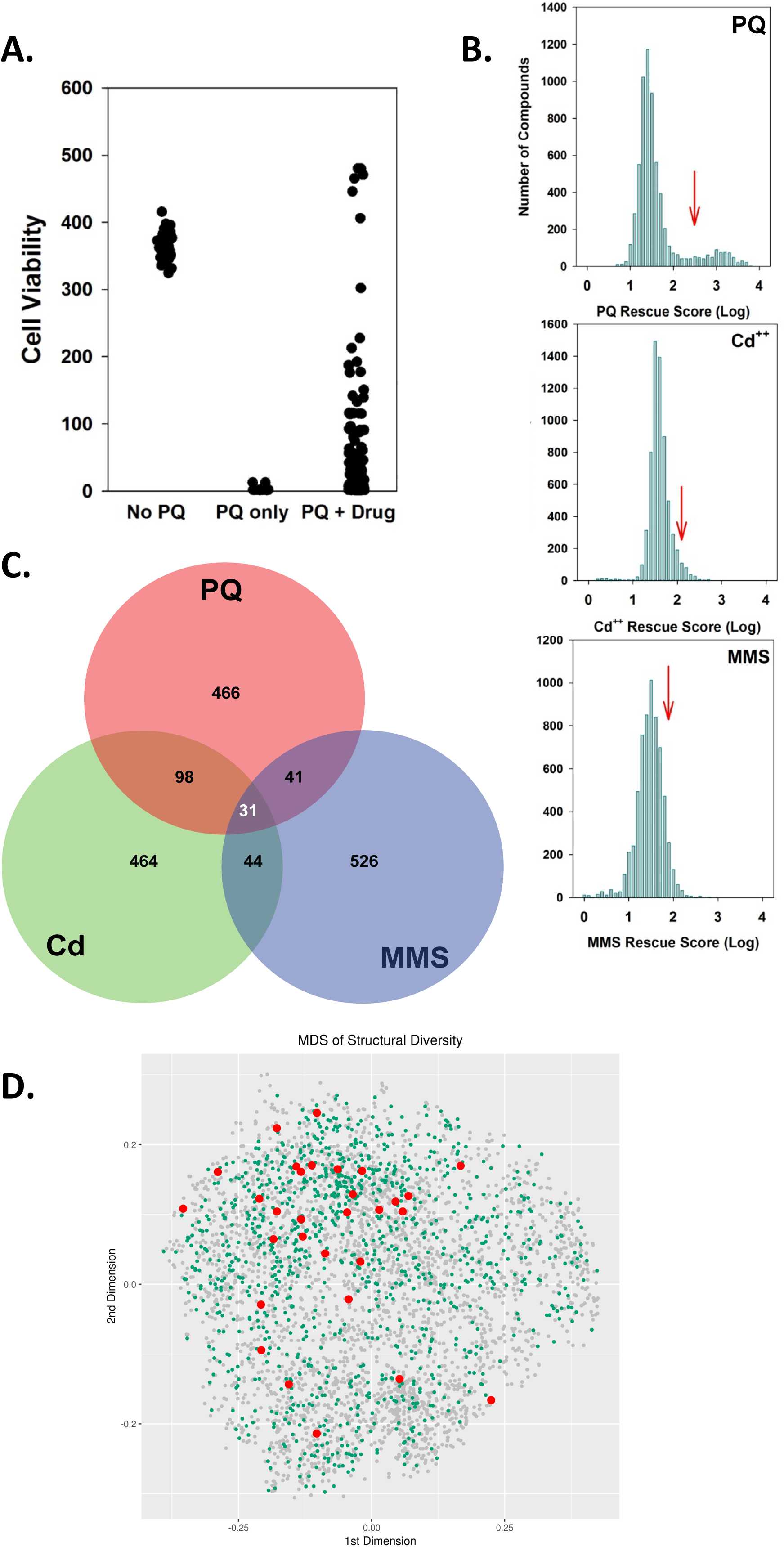
Primary screen for protection of mouse cells from paraquat (PQ) lethality. (A) Example of a single experiment screening 640 compounds. No PQ wells (N=32) received neither PQ nor rescue agent, showing maximum cell viability. PQ only wells (N = 32) show low viability in wells receiving only PQ. PQ + Drug (N = 640) received PQ plus a test agent. Each symbol designates a different well. (B) Histograms for tested agents (N = 6,351), showing number of tested compounds providing specific degrees of cellular protection for PQ, Cd, and MMS. Scores have been normalized by adjustment for plate-to-plate variation in median cell viability, and are expressed on a Log10 scale. Red arrows designate the 90th percentile for each distribution. (C) Proportional Venn diagram showing overlap among compounds in the top 10% for induction of PQ, Cd, and MMS resistance in primary screening. Figure generated using http://www.biovenn.nl/venndiagram.tk/create.php. (D) MDS analysis generated by all-against-all similarity comparisons using atom pairs as structural descriptors and the Tanimoto coefficient as similarity metric. Active compounds in the top 10% for PQ, MMS, and Cd are highlighted as red dots. Compounds active in any of these three stress treatments are highlighted in green.

To facilitate comparisons between experiments and among the different stressors, we calculated an adjusted rescue score for each compound, compensating for minor batch variation so that each test day’s data set shared a common median value (Table S1). Histograms for each stress agent are shown (Fig. 1B), pooling results across four separate experimental batches in each case. Striking differences were observed in the distribution of compounds inducing resistance (Fig. 1B). The distribution of viability scores for PQ was bimodal, with at least 10% of the tested agents promoting PQ resistance at a level 10-fold above the median value of the remainder of the wells (Fig. 1B, top panel, red arrow). In this regard, a prior *C. elegans* RNAi screen for genes regulating PQ resistance showed a similar hit rate as our small molecule screen^32^; thus this relatively high hit rate may reflect the multiple pathways deployed by cells to guard against oxidative insult. In contrast, the histogram for protection against Cd lethality was asymmetric, but not bimodal, with a tail of compounds yielding cell viability scores above the negative control values or the median of the set of tested compounds (Fig. 1B, middle panel). In contrast to the other two stressors, few small molecules induced robust protection against MMS in MTFs under these screening conditions (Fig. 1B, bottom panel).

Compounds that lead to protection against two or more forms of stress (*i.e.* multiplex stress resistance) may be of particular interest from the standpoint of aging biology. Longevity mutations in *C. elegans* often confer resistance to multiple stressors^31^, and MTFs from long-lived mice, such as the Snell, Ames, and GHRKO strains, are also resistant to multiple stresses^23^. To test whether drugs that confer resistance to PQ tend also to impart resistance to Cd and/or MMS, we arbitrarily defined each tested agent as protective for the stress if it produced a cell viability score in the upper 10% of the corresponding distribution (Fig. 1C). Of the 636 compounds in the highest 10% for PQ protection, 129 were also in the highest 10% for Cd protection, significantly higher than the 10% expected by chance (p<10^-6^, permutation testing). There was no statistically significant enrichment among the top 10% of compounds between PQ and MMS, or Cd and MMS. However, among 6,351 compounds tested in the primary screen, 31 (0.49%) conferred at least partial protection against PQ, Cd, and MMS, much more commonly than expected by chance (p<10^-6^, permutation testing). In summary, the majority of compounds that imparted resistance to a stressor in primary screening induced resistance to a single agent only, but multiplex stress resistance occurred more frequently than expected by chance for PQ and Cd, and for all three stressors.

To assess the structural diversity of all screened compounds, multidimensional scaling (MDS) was performed. The distance matrix required for MDS was generated by all-against-all similarity comparisons using atom pairs as structural descriptors and the Tanimoto coefficient as similarity metric (Fig. 1D). The active compounds in the top 10% for all three forms of stress, i.e. PQ, MMS, and Cd in primary screening, are highlighted in red. This analysis emphasizes that compounds that promote multiplex stress resistance are chemically very diverse, and do not closely cluster from a structural perspective.

### Replication and dose-response experiments

We selected 75 compounds from the primary screen results for further study, either because they provided exceptionally strong protection against PQ, or because they were in the top 10% for PQ protection and also provided protection, in the top 10%, for either Cd, MMS, or both (Table S2). Some agents which met these criteria were omitted from secondary dose-response (DR) screening because they were not available in pure form in adequate amounts and/or at reasonable cost. We purchased fresh powders of the chemicals that had been tested in primary screening in the HTS test libraries, and evaluated them over a range of doses (0.3 to 32 μM) for protection against each of the three stress agents. We classified an agent as protective for PQ stress if, in DR testing, at least one dose led to a rescue score of ≥100. For comparison, wells without test agents had a mean rescue score of 10, and wells with no PQ added had a mean rescue score of 820. Thus, categorization as a protective agent required a rescue score 10-fold above the mean negative control. Similar criteria were used to classify the test compounds with respect to protection against Cd or MMS toxicity. This classification was more stringent than that used in the primary screen. Eight of the compounds analyzed in this manner failed DR testing for technical reasons, and were not further considered. Among the 67 chemicals for which we successfully obtained DR curves, 52 (78%) were protective against PQ. Among these, 8 also provided protection against both Cd and MMS, 7 provided protection against Cd but not MMS, 10 protected against MMS but not Cd, and the remaining 16 protected against PQ stress alone. Eleven compounds provided protection against PQ, but failed testing for the other two stressors for technical reasons. Several agents showed a distinct profile of protection in the primary screen versus DR testing. These discrepancies likely reflect the additional data obtained from DR testing, as well as impurities and/or degradation of the samples in libraries used for primary screening.

The DR data for these 67 agents were used to calculate two parameters for each compound: Max, the highest level of protection produced at any dose, and ED50, the lowest concentration that yielded protection at least as high as Max/2. In addition, a drug was classified as toxic if, at the highest tested dose (32 μM), cell viability was less than Max/2. Rescue agents showed diverse patterns in DR testing; representative DR curves are shown (Fig. 2A). For example, dihydrorotenone provided protection at all tested doses. In contrast, supercinnamaldehyde provided substantial protection against PQ at low doses, but was toxic at doses ≥16 μM. Compounds MG 624 and 3230-2939 showed sigmoidal DR curves. ED50 and Max values for each of the tested agents for protection from PQ toxicity are shown (Fig. 2B). Compounds with Max < 100 were considered non-protective (ED50 > 32 μM). Eighteen of the 67 small molecules tested were quite potent, with ED50 values ≤0.5 μM. The maximum level of protection did not correlate with ED50 in this set of tested agents.

Max and ED50 values, for PQ, Cd, and MMS for each of the 67 compounds with DR data are provided (Table S2). An overview of relationships between Max values for PQ and Max values for Cd for each drug is shown (Fig. 2C). Many agents provided robust protection against PQ but not Cd, and a few protected against Cd but not PQ, despite having been initially selected based on a screen for PQ resistance at the single 16 μM dose in primary screening. Drugs that yielded protection against MMS are indicated (Fig. 2C, blue triangles). Seven compounds conveyed Max > 100 for all three stress agents. However, there was not a strong correlation between Max values for PQ and Cd, nor a strong tendency for agents that protect against PQ and Cd to protect cells against MMS as well.

**Figure 2:**
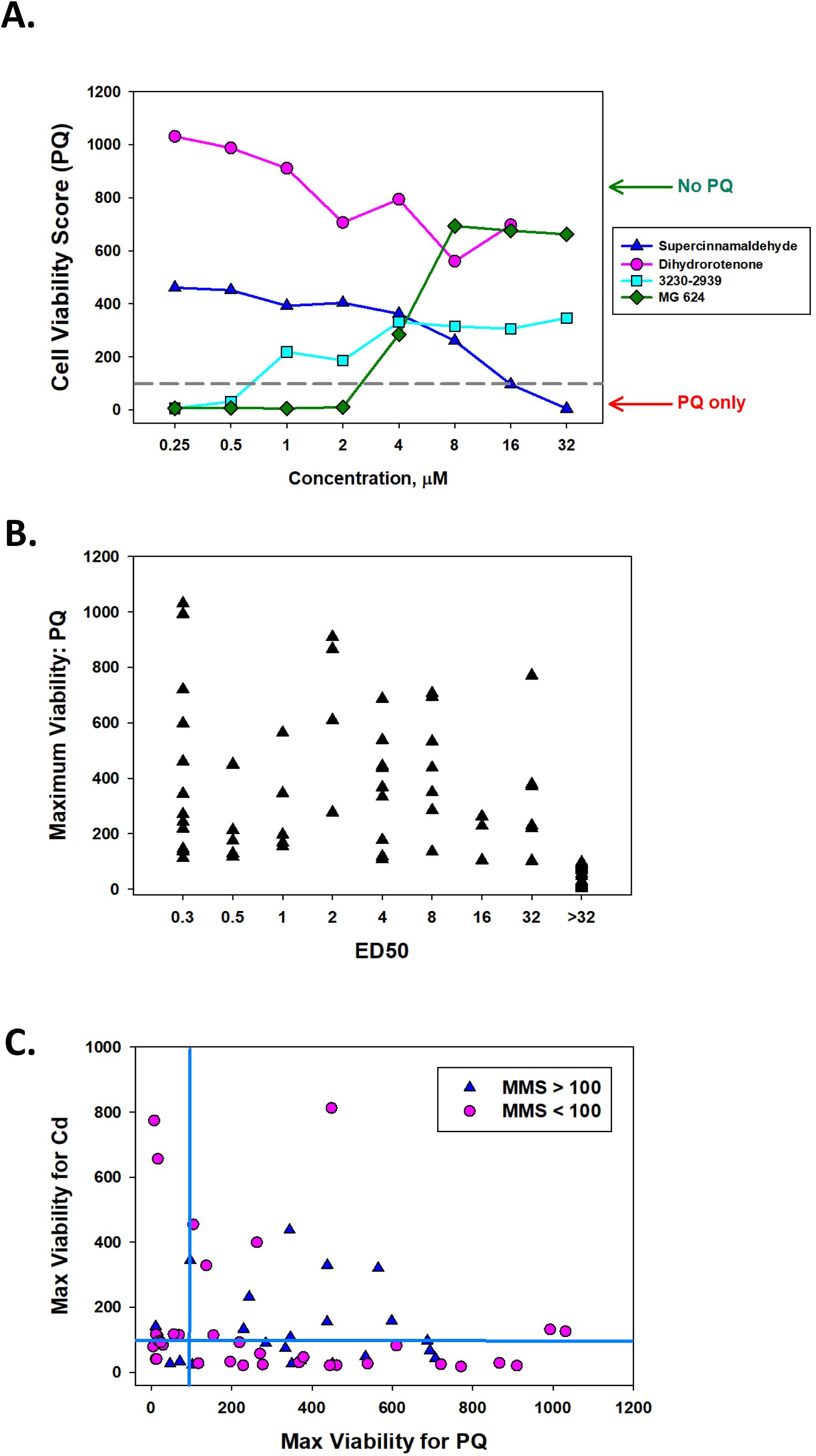
Dose-response curves for selected small molecules. (A) Representative dose-response curves, for concentrations from 32 to 0.25 μM (X axis). Green arrow designates mean score (from four daily batches) for wells with neither PQ nor protective agent added, and red arrow designates mean score for wells that received PQ alone. Grey dashed line shows a viability score of 100, used as an arbitrary criterion for categorizing an agent as protective. (B) Scatterplot for ED50 vs Max viability score for PQ protection; each symbol represents a different rescue agent. (C) Scatterplot showing Max score for PQ protection versus Max protection for Cd stress. Blue triangles indicate rescue agents for which Max > 100 in the MMS protection dose testing. Dashed lines indicate arbitrary thresholds at Max = 100 for PQ and Cd protection scores.

### Relationship between induction of MTF stress resistance by small molecules and lifespan extension in *Drosophila* or *C. elegans*

Since enhanced stress resistance is a conserved feature of extended longevity, we tested whether compounds identified in HTS in MTFs would extend lifespan in *Drosophila melanogaster* and *C. elegans,* invertebrate models often used for studies of aging and lifespan. We fed both *C. elegans* and *D. melanogaster* selected small molecules over the entire lifespan and assayed survivorship (Table S3), to test the hypothesis that compounds that improved cellular survival in response to stress would be more likely to induce invertebrate lifespan extension compared to compounds with little or no effect on MTF survival.

In flies, drugs were tested at 20 μM in females, and at 200 μM in males. A compound was scored as positive if it increased lifespan in either sex compared to DMSO control, by log-rank test at p=0.05 or better. Among the 70 unique compounds that significantly increased MTF stress resistance that were tested in flies, 12 significantly increased fly lifespan in either or both sexes (17.1% of the total compounds tested) (Table S3). However, 16/54 negative control compounds tested (*i.e.*, those that did not induce MTF stress resistance) were also able to significantly increase fly lifespan (29.6% of the total compounds tested; p=0.07 for hits versus negative controls). No tested agent increased median lifespan by more than 20%. In *C. elegans,* compounds were tested in hermaphrodite worms at two concentrations, 10 μM and 100 μM (Table S3). A compound was scored as positive if it significantly increased lifespan by log-rank test at p=0.05 or better, at one or both doses. 15/70 (21.4%) compounds that increased MTF stress resistance extended worm lifespan, as did 6/54 (11.1%) negative control compounds (p=0.15). If we consider compounds that extended lifespan in either or both invertebrate organisms, 26/70 hit compounds extended lifespan (37.0%), as did 20/54 negative control compounds (p=0.99). Thus, in this relatively small dataset, a set of agents selected for their ability to protect MTFs from stress were not enriched in compounds that extend fly or worm lifespan.

To extend this analysis, we took advantage of the fact that one of the libraries screened for lifespan effects in our primary MTF screen, the LOPAC collection, had previously been assayed for effects on worm lifespan^18^. Among the 1,280 compounds in the LOPAC library, we were able to match 827 between our HTS and the data tabulated by Ye *et al*. A caveat to this analysis is that, since the latter publication provided no compound structure information, matching entries among the two data sets used chemical names, and formatting differences may have introduced errors. We considered “hits” from the HTS to be those that were in the top 10% for PQ, MMS, or Cd; this amounted to 207/827, or 25% of the total (Table S4). We considered the “hits” from the Ye paper to be compounds in the top 10% of positive change in lifespan, representing a 15% lifespan increase or better. Among the 207 hits from our HTS, 34 were also in the top 10% for lifespan increase (16.4%). However, among the 620 compounds in the bottom 90% for all three stressors in the cell screen, only 40 were in the top 10% for increased worm lifespan in the results of Ye and colleagues (6.5%), indicating a highly significant association between induction of cellular stress resistance and *C. elegans* longevity (p<0.0001, Fisher’s exact test). In summary, our own small-scale analysis did not identify a significant relationship between small molecule induction MTF stress resistance and lifespan extension in *C. elegans*, although better-powered comparison with a published *C. elegans* screen did suggest such a relationship.

Interestingly, in our *C. elegans* lifespan studies, among the agents scoring positive for extension of lifespan at both 10 and 100 mM, in two separate trials, was the antibiotic tetracycline (Fig. S2A). This is consistent with prior results showing that interventions that selectively impair mitochondrial protein synthesis can increased worm lifespan^39^. Likewise, the mitochondrial poison dihydrorotenone increased worm lifespan at two doses (Fig. S2B), consistent with the known ability of mitochondrial defects present at specific developmental stages to extend longevity in this organism^40, 41^. Both of these agents robustly promoted PQ resistance in MTFs (Table S1), suggesting that these drugs may exert conserved effects in worms and mammalian cells, potentially via impairment of mitochondrial function.

### Associating screening hits with known drugs, target proteins and pathways

We used fragment-based structural similarity searches combined with the Tanimoto coefficient to identify nearest structural neighbors in DrugBank^42^ for the 4,920 unique agents for which we had acquired cell stress resistance data in primary screening^43, 44^. DrugBank was chosen as the reference database for this purpose because it represents one of the most comprehensive and best curated collections of drugs available in the public domain. This allowed us to obtain detailed functional annotations from DrugBank for many of our screened compounds, including therapeutic usage and target proteins^45, 46^. Among the compounds used for our primary screens, 1,935 (39%) were represented in DrugBank based on a Tanimoto coefficient of ≥0.9. Nearly all DrugBank compounds for our screening hits were annotated with at least one target protein. The results of this in silico analysis, including target protein annotations, are presented in Supplementary Tables S5 and S6.

We then performed functional enrichment analysis (FEA) on the groups of compounds identified as hits in the PQ and Cd screens, as well as both of them (PQ and Cd). As functional annotations systems we included here Gene Ontologies (GO), Disease Ontologies (DO) or KEGG pathways^47–50^. Two complementary enrichment methods were used for this: Target Set Enrichment Analysis (TSEA) and Drug Set Enrichment Analysis (DSEA). For TSEA the compound sets were converted into target protein sets based on the drug-target annotations obtained from DrugBank in the previous step. The corresponding gene sets for these target sets were used as test samples to perform TSEA with the hypergeometric distribution. Since several compounds in a test set may bind to the same target, the test sets can contain duplicate entries. This corresponding frequency information will be lost in traditional enrichment tests, because they assume uniqueness in their test sets, which is undesirable for FEA of compounds. To overcome this limitation, we also performed DSEA on the active compound sets themselves (here test sets) against a database containing compound-to-functional category mappings. The latter was generated by substituting the targets in the former by the compounds they bind. The main advantage of DSEA is that it maintains functional enrichment information in situations when several compounds bind to the same target, because there are usually no duplicates in compound test sets. The enrichment results for TSEA and DSEA are available in Supplementary Tables S7 and S8, respectively.

As expected, the DSEA results are not identical to the TSEA results, but share several top ranking functional categories. For brevity and consistency, the following evaluation of the FEA will focus on the TSEA results. Interestingly, the most highly enriched functional categories found in the TSEA for the PQ (Fig. 3B) and Cd (Fig. 3C) screens were very similar with respect to both their compositions as well as their rankings by enrichment p-values. The most common GO term and KEGG pathway annotations among the top scoring categories are related to receptor and transporter functions, as well as signaling pathways. Another interesting finding is that the two most highly enriched DO terms in both the PQ and Cd screen (Supplementary Table S7) are associated with age related diseases, such as congestive heart failure (DOID:6000; adjusted p-value=2.08×10^-6^) and heart disease (DOID:114, adjusted p-value=2.35×10^-6^). Based on the pharmacodynamics annotations from DrugBank (Supplementary Table S5), the test sets used for the enrichment analyses of both screens include FDA approved drugs that are used to treat cardiovascular conditions.

**Figure 3:**
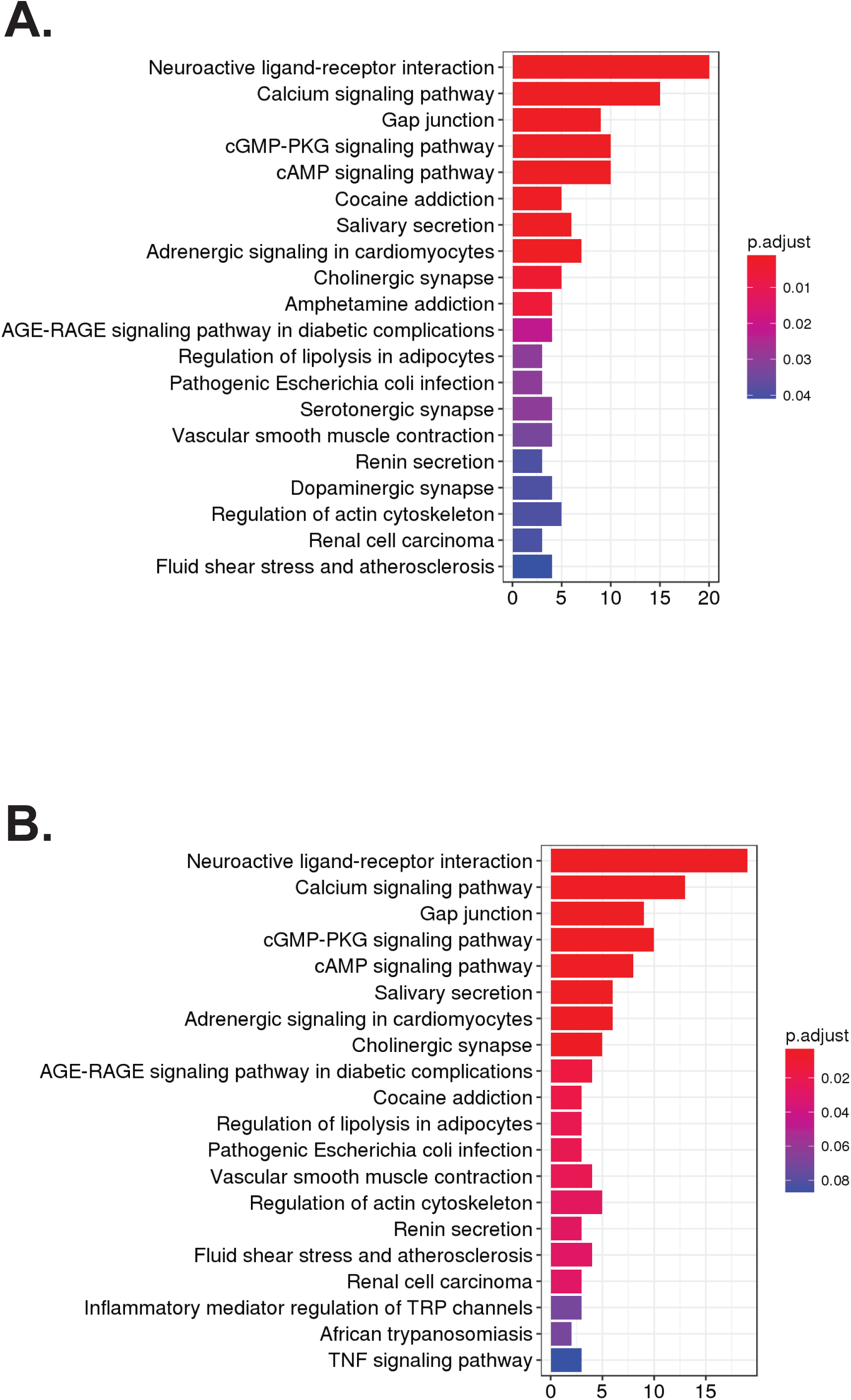
Chemoinformatic analysis of small molecules that promote stress resistance. Drug set enrichment analysis (DSEA) with KEGG pathways for PQ (A) and Cd (B). Drugs where associated with KEGG pathways based on drug-target annotations of their structural nearest neighbors in DrugBank.

### Transcriptomic analysis show that AEG3482 and Cardamonin activate Nrf2 signaling

To gain additional mechanistic insight into how compounds identified by HTS confer stress resistance, we performed RNA-seq analysis on MTFs treated with 8 compounds: AEG 3482, antimycin A, berberine HCl, cardamonin, clofilium tosylate, dihydrorotenone, diphenyleneiondonium HCl, and podofilox^51–53^. These compounds were selected due to their robust induction of stress resistance, modest cost, and biological interest. Principal component analysis (PCA) of the RNA-seq data plotted on the first two principal components showed that 5 compounds clustered together with DMSO in PC1, and to a lesser extent, in PC2 (Fig. 4A). In contrast, three other compounds (AEG 3482, cardamonin, and podofilox) induced a much greater degree of transcriptomic heterogeneity, and differed from one another markedly in PC1. Cardamonin and podofilox, but not AEG 3482, had similar values for PC2 (Fig. 4A).

**Figure 4.**
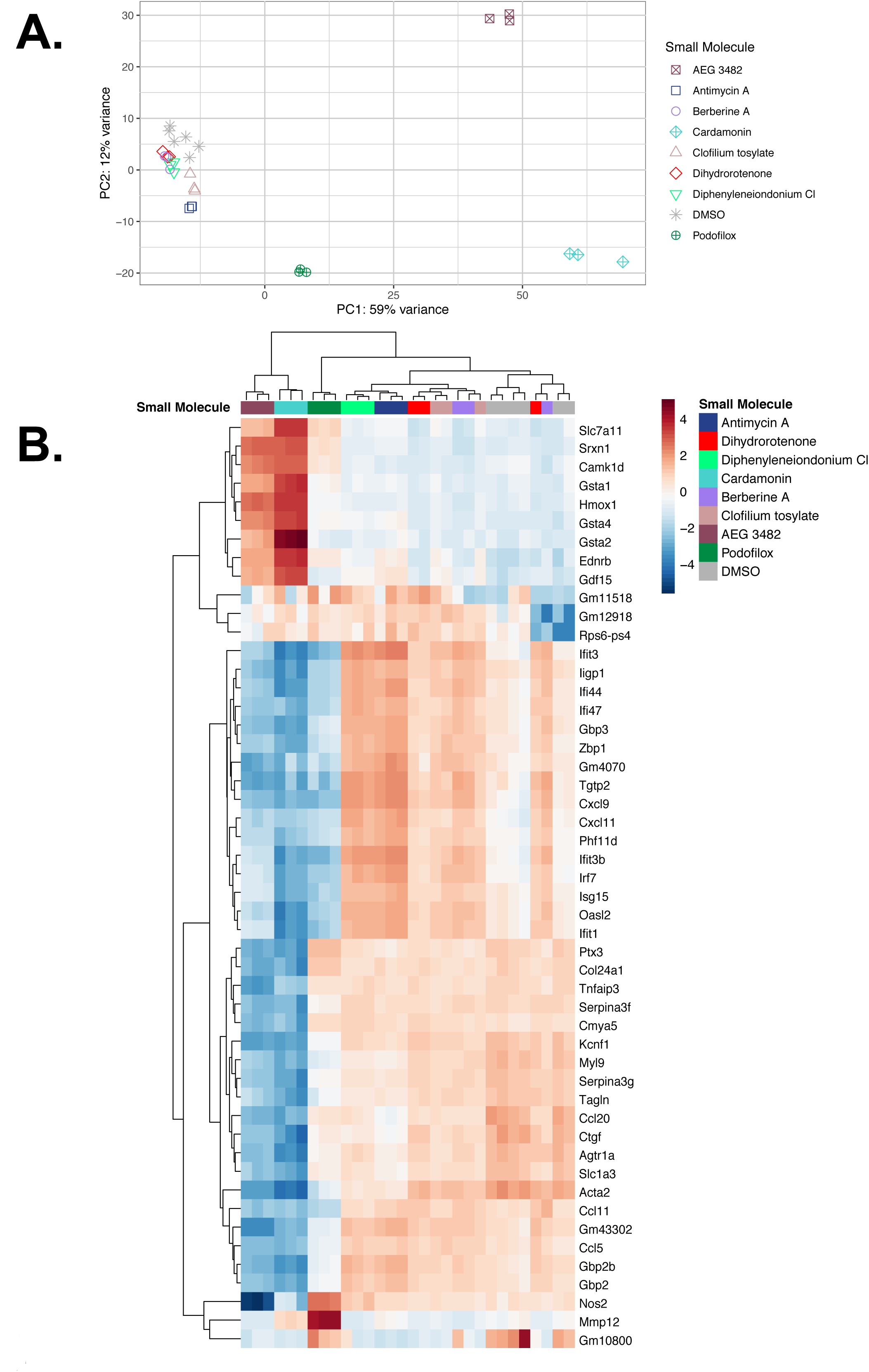
Transcriptomic analysis of eight rescue compounds. (A) Principle component analysis (PCA) for all RNA-seq samples. (B) Dendrogram with unsupervised hierarchal clustering of the top 50 genes with greatest variance from the mean. Expression across each gene (row) has been scaled such that the mean expression is zero (pale yellow). Red indicates high expression in comparison to the mean. Blue indicates low expression in comparison to the mean. Lighter shades represent intermediate levels of expression.

To identify differentially expressed (DE) genes for each compound compared to the DMSO control, a significance threshold of FDR < 0.05 and log_2_fold-change threshold of >1 or < -1 were imposed. Unsupervised hierarchical clustering analysis (HCA) based on the top 50 genes with greatest variance from the mean revealed that AEG 3482 and cardamonin clustered together, while the other six small molecules tested clustered together with each other and with DMSO control (Fig. 4B). In analysis of a larger group of the 300 most variant genes, this clustering was maintained (Fig. S3). Overall, cardomonin and AEG 3482 induced (1) a larger number of differentially expressed (DE) transcripts, and (2) changes in these transcripts of greater magnitude compared to DMSO control (Table S9).

To elucidate potential mechanisms by which these compounds induce stress resistance, we performed functional analysis of DE genes using Qiagen Ingenuity Pathways Analysis (IPA) and gene set enrichment analysis (GSEA)^54^. Interestingly, cellular immune response and cytokine signaling-related terms were identified as significantly affected IPA-defined pathways for all 8 compounds; many of the genes involved relate to cytokine signaling and innate immunity (Table S10). For AEG 3482 and cardamonin, manual inspection of IPA and GSEA results revealed a strong oxidative stress resistance signature, including NRF2-mediated oxidative stress response (Table S11). This strong NRF2 antioxidant gene expression signature was not evident for the six other compounds (Table S11).

The transcription factor nuclear factor erythroid 2-related factor 2 (NFE2L2, aka Nrf2) is a core component of the cellular response to oxidative stress^55^. IPA revealed several pathways related to Nrf2 oxidative stress induced by AEG 3482 and cardamonin (Fig. 5A). Likewise, according to GSEA, Nrf2 induction and a key transcriptional target pathway of Nrf2, Glutathione Conjugation, were among the most highly enriched categories for cardamonin (Fig. 5B) and AEG 3482 (Fig. S4). Glutathione S-transferases neutralize a wide range of electrophilic molecules, including reactive oxygen species (ROS)^55^. Other antioxidant defense mechanisms regulated by Nrf2 include synthesis and regeneration of cellular reducing agents – *i.e.* GSH – expression of genes in the antioxidant thioredoxin (TXN) system, and regulation of heme metabolism^56^.

**Figure 5:**
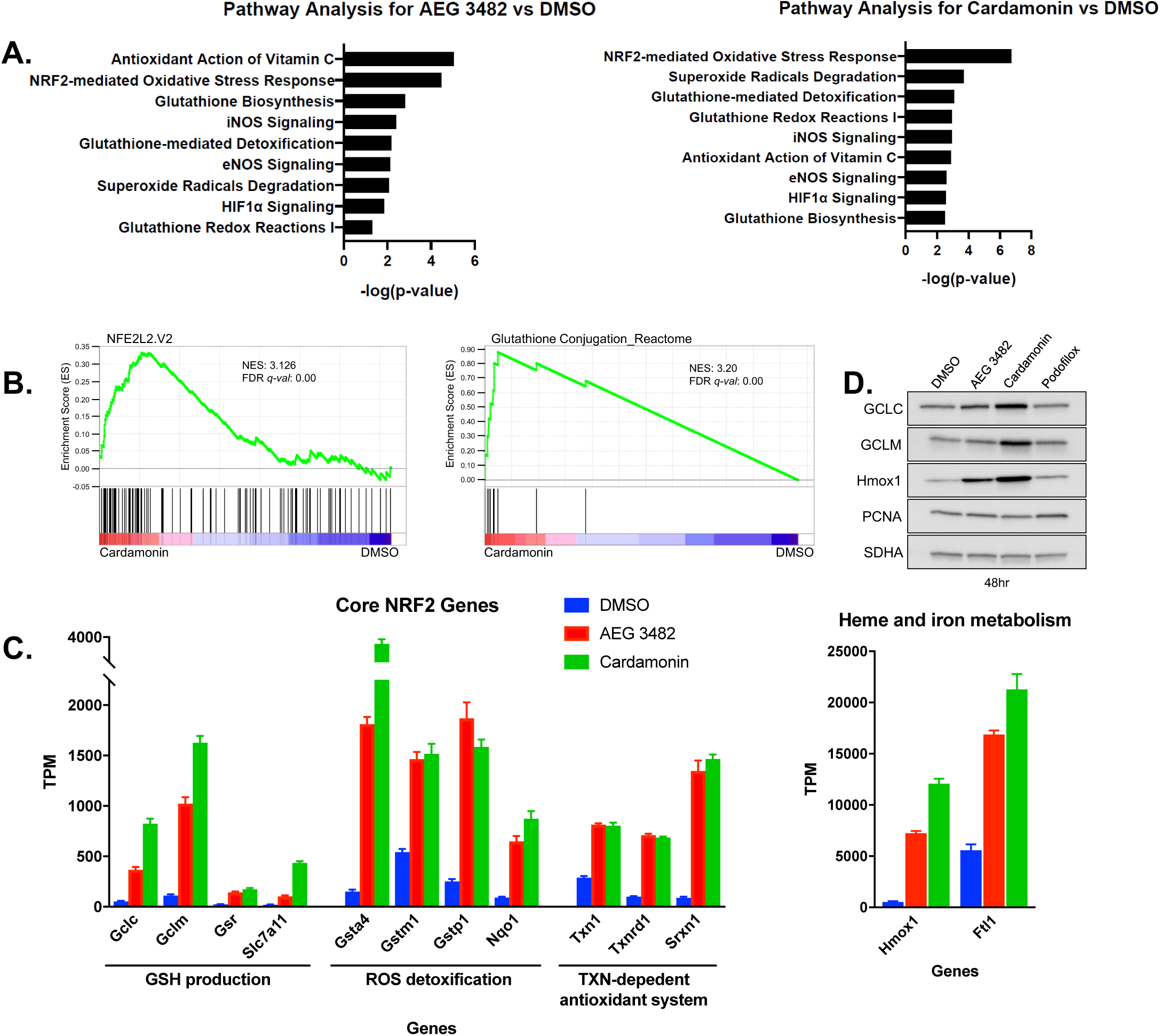
Nrf2 antioxidant signature in response to AEG 3482 or cardamonin treatment. (A) Pathways related to oxidative stress response from IPA for DE genes induced by AEG 3482 and cardominin. P-values represented in negative log form; all pathways are significant at p<0.05. (B) Gene set enrichment analysis (GSEA) for cardamonin for pathways indicated. (C) Transcripts in key pathways induced by Nrf2 are significantly elevated by AEG 3482 and cardamonin treatment. All false discovery rates (FDR) for comparison to DMSO are q < 5*10^-6^. (D) Immunoblot analysis of Nrf2 targets after treatment with indicated compounds for 48 hours. PCNA and SDHA serve as loading controls.

Under basal conditions, Nrf2 is actively polyubiquitinated and degraded by the Keap1/Cul3 complex, but is stabilized in response to various insults, in particular oxidative or xenobiotic stress^55^. In RNA-seq analysis, *NFE2L2* mRNA was not induced by any of the compounds when compared to the DMSO control. From the RNA-seq analysis, we also extracted the best-characterized genes representative of a Nrf2 response signature (Fig. 5C). Notably, expression of genes involved in GSH synthesis, utilization, and regeneration were all dramatically increased in response to AEG 3482 and cardamonin. In addition to GSH-related genes, AEG 3482 and cardamonin also induced expression of genes in the thioredoxin (TXN) pathway, which reduces oxidized protein thiols (Fig. 5C). These changes did not occur in response to the 6 other compounds tested (Fig. S5). Oxidized TXN is reduced to a functional form by thioredoxin reductase (*Txnrd1*)^57^. Transcripts encoding heme oxygenase (Hmox1) and Ftl1, important for the proper catalytic degradation of heme and Fe^2+^ storage, were also up-regulated by AEG 3482 and cardamonin (Fig. 5C). Improper degradation of heme can result in generation of free Fe^2+^, which can then catalyze conversion of H_2_O_2_ to damaging hydroxide radical^56^. To confirm that changes observed by RNA-seq were reflected at the protein level, we performed immunoblot analysis for GCLC and GCLM (subunits of glutamate cysteine ligase, involved in the glutathione synthesis pathway; Fig. 5D), and Hmox1 at 48 hrs after compound treatment. Consistent with mRNA expression data, AEG 3482 and cardamonin, but not podofilox, increased protein expression of these Nrf2 targets at 48 h (Fig. 5D).

### Proteomic analysis reveals widespread changes in cellular signaling pathways induced by hit compounds

Besides AEG 3482 and cardamonin, the other six compounds did not induce a Nrf2 transcriptional signature, suggesting that they function through other pathways to promote cellular stress resistance. To identify such candidate pathways, we performed Reverse Phase Protein Array (RRPA) analysis on the same eight compounds tested in RNA-seq. RPPA is a high-throughput antibody-based approach that permits simultaneous assessment of levels of hundreds of proteins and phosphoproteins^58^. In this analysis, we found that the eight compounds induced apparent changes in multiple signaling pathways relevant for cellular stress resistance, survival, and longevity, including AMPK, AKT, mTOR, and others (Figs. 6A and S6).

We focused on validating changes in one of these pathways, AMPK signaling. AMPK is a heterotrimeric kinase involved in orchestrating cellular responses to reduced energy charge that also plays roles in cellular stress responses, via regulation of autophagy and apoptotic signaling^59^. Phosphorylation of Threonine 172 (T172) on the α subunit of AMPK is required for AMPK activation. RPPA analysis showed that cell treatment with diphenyleneiondonium HCl, berberine HCl, antimycin A, or clofilium tosylate all significantly increased AMPK T172 phosphorylation (Fig. 6A and Table S12). Conversely, cardamonin, AEG 3482, dihydrorotenone, and podofilox reduced p-AMPK. For all compounds tested except clofilium and diphenyleneiondonium, similar changes were observed for p-ACC1, a major downstream target of AMPK signaling. We were able to validate some of these effects via direct immunoblot; AEG 3482 significantly reduced p-AMPK levels, whereas antimycin, berberine, and clofilium all significantly increased it (Fig. 6B-C). We conclude that specific hit compounds from our screen can modulate signaling through AMPK, and likely other cellular pathways important in stress resistance, potentially helping to explain their effects on cell survival.

**Figure 6:**
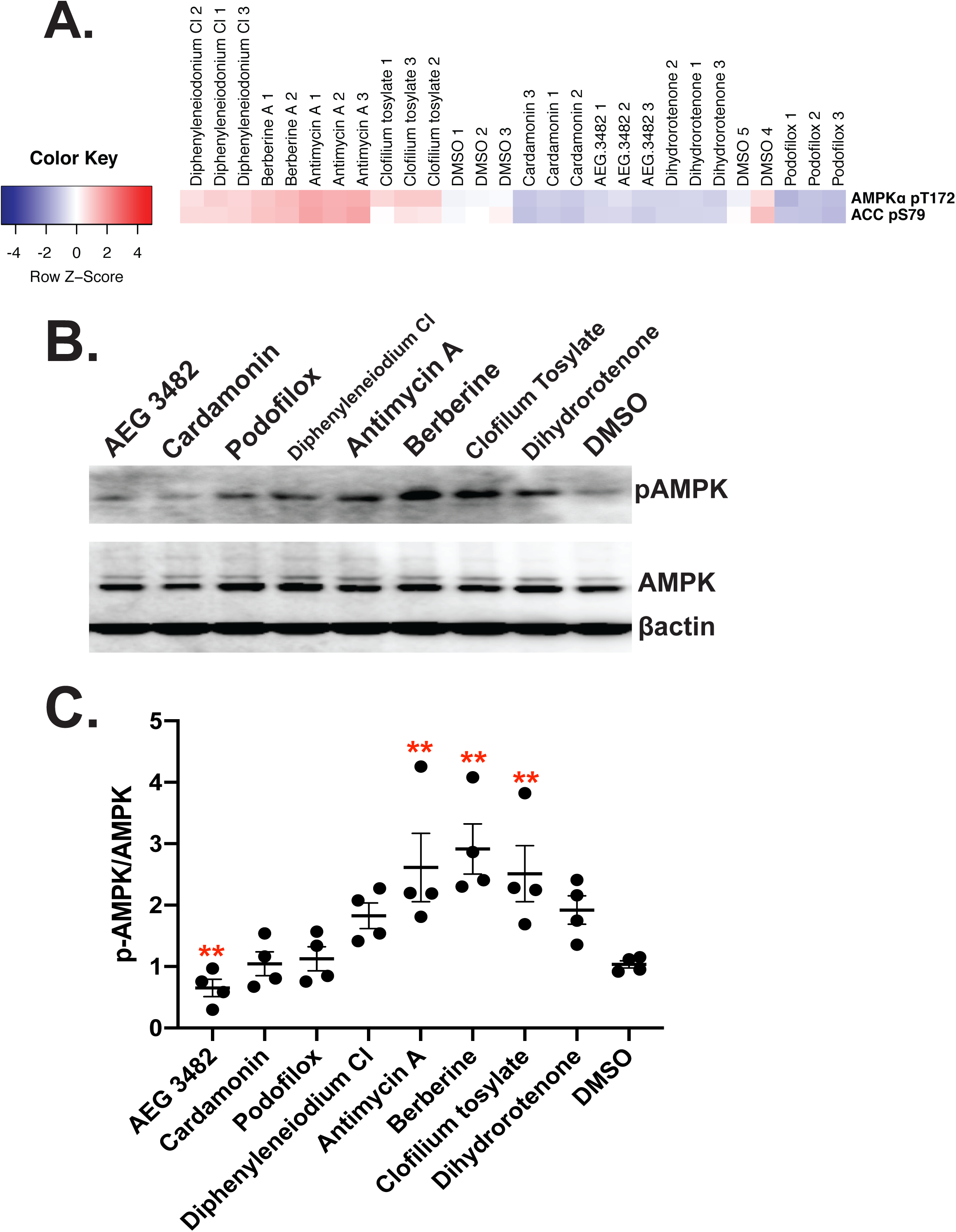
Hits from HTS affect signaling through AMPK and other cellular pathways. (A) RPPA results for p-AMPK and p-ACC1 for 8 compounds tested. (B) Representative immunoblots showing effects of treatment with the agents indicated at 4 μM for 24 hours. (C) Summary of four studies similar to (B). Ratios of p-AMPK/AMPK relative to untreated control are shown; ** p<0.01 (Student’s two-tailed t-test).

## Discussion

The unbiased identification of compounds that can enhance mammalian health- and lifespan represents a major challenge. Resistance to multiple forms of environmental insult is a common feature of longevity, among different species, in invertebrate aging models, and in genetically engineered strains of mice^23^. In this work, we performed HTS to identify small molecules that promote stress resistance in mouse fibroblasts, and identified compounds promoting survival in response to PQ, Cd, MMS, as well as multiplex stress resistance. The ability of small molecules to increase fibroblast stress resistance was not strongly correlated with extension of worm or fly lifespan in our own smaller-scale experiments, but there was a strong correlation in a larger, previously published dataset^18^. Transcriptomic analysis identified Nrf2 signaling as a likely pathway involved in stress resistance provoked by two hits, cardamonin and AEG 3482. Some of the compounds identified in our studies – particularly those that induce multiplex stress resistance – may merit further mechanistic studies, and testing in mice for their ability to increase healthy lifespan.

Some of the compounds identified as hits in our HTS have previously been investigated for their potential health benefits. For example, epigallocatechin-3-monogallate (EGCG) – which promotes PQ and MMS resistance in our DR studies – is a polyphenol present in green tea, purported to possess beneficial properties against cancer and cardiovascular disease^60^. Carvedilol, found to promote PQ and Cd resistance in our DR studies, is a non-selective β-adrenergic blocker with anti-α-1 activity, clinically useful in studies, is a non-selective patients with heart failure^61^. It is possible that some of the cardioprotective effects of this agent occur in part through promotion of cellular stress resistance in cardiomyocytes, although importantly we have not directly assessed the effects of carvedilol on this cell type. Likewise, cardamonin is a spice-derived nutraceutical with anti-neoplastic and anti-inflammatory properties, which may involve modulation of the activity of STAT3 and other transcription factors^62^.

It is notable that several compounds that induce PQ resistance – antimycin A, dihydrorotenone, rotenone, tetracycline, chloramphenicol, and rifampicin (Table S1) – are known mitochondrial toxins. Cellular PQ injury arises through redox cycling, whereby PQ is reduced in an NADPH-dependent reaction, in turn transferring an electron to O_2_ to form toxic superoxide. The mitochondrion represents an important site for PQ reduction^63^; hence interference with mitochondrial function by these mitochondrial toxins identified in our screen may impair the ability of this organelle to cycle PQ. Alternatively, or in addition, this phenomenon may reflect hormesis^64^: *i.e.* the cellular response to mitochondrial dysfunction induced by these rescue agents may prime the cell to more effectively respond to PQ toxicity.

Our approach has several potential limitations. Cellular stress resistance is a complex phenotype influenced by numerous signaling pathways, which may have led to the surprisingly high initial hit rate (approximately 10% against PQ in primary screening), and the chemical diversity and large number of potential targets of our protective compounds. In this regard, it should be noted that an RNAi screen in *C. elegans* for genes regulating PQ resistance showed a similar hit rate as our small molecule screen^32^. Our primary screening was carried out at a single concentration of rescue compound (16 μM), thus potentially missing chemicals that may be effective at lower doses, but ineffective or toxic at the particular dose used in the screen. Compounds that increase stress resistance of specific cell types, such as neurons or hepatocytes, might be missed by our screening strategy, since we have focused on only a single cell type, MTFs. Similarly, our strategy will not identify compounds that require metabolic transformation *in vivo*, *e.g.* in the liver, for biological activity.

Compounds that increased resistance of mouse fibroblasts to multiple forms of stress were enriched, compared to the overall set of tested agents, for the ability to promote *C. elegans* lifespan in a prior screen^18^. However no such relationship existed in our smaller-scale *C. elegans* and *Drosophila* studies. There are many caveats to the invertebrate experiments. For example, we do not know whether worms and flies consume food similarly on all compounds administered to them, nor do we know the optimal dose of each compound in the invertebrate models, nor how well the compounds may penetrate the cuticle. Overall the data suggest that, as a general matter, it is not straightforward to identify compounds that increase invertebrate lifespan by selecting drugs that increase stress resistance of cultured mammalian cells, unless a large number of compounds are screened. Nevertheless, PQ resistance has successfully been used as a surrogate phenotype in an RNAi screen to identify longevity mutants in *C. elegans*^32^. Moreover, in worms, many mutants with increased lifespan also show elevated stress resistance^31^. Notable among these is the *daf-*2 mutant, which displays greatly extended longevity as well as multiplex stress resistance: enhanced survival in response to oxidative, heavy metal, heat, and infectious insults. These findings emphasize the evolutionarily conserved nature of the link between oxidative stress resistance and lifespan.

Among the 8 compounds analyzed via RNA-seq, two – cardamonin and AEG 3482 – strongly activated Nrf2 signaling and antioxidant defenses, potentially accounting for their ability to render cells stress-resistant. Nrf2 regulates expression of numerous cytoprotective genes via binding to promoter antioxidant response elements (AREs) in response to oxidative or xenobiotic insult. Nrf2 inducers are under development for a variety of inflammatory, metabolic, degenerative, and neoplastic diseases^65^. The induction of Nrf2 signaling by cardamonin that we observe is consistent with prior reports in the literature^66–68^. In this regard, NRF2 levels and target gene expression are elevated in skin fibroblasts derived from long-lived Snell dwarf mice, likely contributing to the multiplex stress resistance observed in these cells^37^. Similarly, Protandim, a mixture of botanical extracts that induces Nrf2, extends median lifespan in male mice^69^. These observations support the notion that some of the hits identified using this HTS approach may exert beneficial longevity effects in mammals.

AEG 3482 was originally identified by its ability to suppress death of cultured neurons following growth factor withdrawal^70^. It is thought to act via binding to HSP90 to induce HSF1 activity, increase HSP70 expression, and suppress JNK signaling. In MTFs, using our experimental conditions, AEG 3482 provoked only a weak “Cellular Response to Heat” signature by GSEA (Fig. S7A); the related signature, “Heat Shock Protein Response”, was non-significantly induced (not shown). Under our treatment conditions, neither AEG 3482 nor cardamonin induced HSP70 or HSP90, whereas podofilox treatment increased HSP90 expression only at a late timepoint (Fig. S7B). Importantly, extensive cross-talk between HSF1 and NRF2 signaling occurs; these transcription factors are known to regulate overlapping sets of genes^71^. Chemically distinct NRF2 activators are reported to induce an HSF1 target gene^72^. Thus, it is possible that HSF1 responses contribute to the stress resistance induced by AEG 3482 and/or cardamonin.

The other six small molecules analyzed did not produce a clear NRF2 or antioxidant gene expression signature. These compounds may promote stress resistance via their effects on cellular inflammatory pathways, or by regulating other upstream signaling pathways. Future studies are needed to elucidate their mechanism of action.

In summary, we have performed HTS to identify compounds that confer protection against multiple forms of cytotoxic insult to mammalian cells. These hits – particularly the agents that induce resistance to multiple stressors – may prove to be useful tools to probe mechanisms of cellular stress resistance, and to test for beneficial effects in rodent models. More broadly, the strategy presented here may represent a workflow for screening larger compound libraries to identify new candidate anti-aging molecules with potential health or lifespan benefits in humans.

## Acknowledgements

This work was supported by the Glenn Foundation for Medical Research. AG was supported by T32 GM113900 and T32 AG000114. The authors would also like to cite the following grants: R01GM101171 and R01HL114858 (Lombard); R01AG030593 and R01AG051649 (Pletcher); U24AG051129, 1S10-OD016290-01A1, and NSF award ABI-1661152 (Girke). The authors thank Dr. Jun Qi (Dana Farber Cancer Institute) for (+)-JQ1 and (-)-JQ1, and Drs. Michael Petrascheck and Matthias Truttmann for helpful discussions. Some studies reported in this publication were carried out at the Center for Chemical Genomics (CCG) at the University of Michigan Life Sciences Institute. RNA-seq results have been deposited at GEO, accession number GSE130294.

## Materials and Methods

### Cell generation

Mouse fibroblast cell lines were established using pooled tail fibroblasts from 5 male and 5 female UMHET3 mice, each 8 to 11 weeks old. Cell lines were started from each donor mouse and grown to a minimum of 20 million cells simultaneously. 20 million cells from each donor mouse were thoroughly mixed and aliquots of 2 × 10^6^ cells were cryopreserved at passage 2. Cells were thawed as needed and passaged twice after thawing. At the second passage cells were suspended in high glucose DMEM without sodium pyruvate and supplemented with 10% fetal bovine serum plus penicillin/streptomycin/Amphotericin B. Cells were kept in a 37°C, 10% CO_2_, humidified incubator.

### Stress resistance assays

One day 1, MTFs were suspended at a concentration of 10,000 cells/25 μl (=0.4×10^6^ cells/ml) in DMEM plus 0.5% BSA. Cells were then aliquoted into 384-well plates (Greiner cat# 781080) and incubated overnight at 37° C in a 10% humified CO_2_ incubator. On day 2, 0.2 μl of test compounds (plate columns 3-22) or DMSO (plate columns 1-2 and 23-24) were added, for an initial compound concentration of 16 μM and initial DMSO concentration of 0.8%. Plates were then incubated overnight. On day 3, 5 μl of media alone (columns 1-2) or media plus stress agent (columns 3-24) were added to plates. Typical final concentrations of stress agent used in primary screens were PQ, 12 mM; Cd, 4 μM; and MMS, 0.76 mM. In some studies, 15 mM PQ and 12 μM Cd were used in parallel with the more standard stress conditions. On day 4, 5 μl of Cell Titer Glow detection reagent was added to each well, and luminescence was measured. The maximal value was defined as the average luminescence value of DMSO-treated cells without stress agent in column 1. The minimal value was defined by the average of the cells from column 24 treated with DMSO with stress agent. The rescue score for each compound tested was defined as (well value – plate average of DMSO+stress agent)/plate average DMSO only value).

### Cheminformatics and Drug-Target Analyses

Reference compound structures were downloaded from DrugBank in SDF format^42^. Analyses of small molecule structures were performed with the *ChemmineR* package^43, 44^. Structure similarity searches of small molecules used atom pairs^73^ as fragment-based descriptors and the Tanimoto coefficient as similarity metric^45, 46^. Analyses of drug-target annotations from DrugBank were performed with the *drugbankR* package (unpublished). Target set and drug set enrichment analyses (TSEA and DSEA) used the hypergeometric distribution test along with the Benjamini & Hochberg (BH) method for adjusting p-values for multiple testing implemented by the *clusterProfiler* package^47^. Custom R functions were used for generating compound-to-KEGG pathway mappings.

### Cell preparation for RNA-seq and RPPA

For RNA-seq, 0.5 × 10^6^ cells were added to each of 4 wells in a 6-well plate per sample. Total volume per well was 3 ml (day 1). Cells were left to adhere overnight. On day 2 the compounds were added to the cells and the cells were left to incubate overnight. On day 3, media was removed, cells were washed two times with PBS. RNazol was then added directly to the cells and collected for RNA seq analysis. For RPPA, 10^6^ cells were added to one T-175 cm^2^ flask for each sample. Total volume per flask was 20 ml (day 1). Cells were left to adhere overnight.

On day 2 the compounds were added to the cells and the cells were left to incubate overnight. On day 3 RPPA samples were collected using 0.05% trypsin, trypsin was neutralized after cells detached by adding complete media, and cells were washed twice with PBS. RPPA was performed at the RPPA Core Facility at MD Anderson Cancer Center. Further information about the compounds used in RNA-seq and RPPA is provided in the supplemental methods, including vendors, catalog numbers, CAS numbers, and SMILES structures.

### RNA-sequencing bioinformatics processing

Reads were aligned to mm10 using kallisto v0.43.1. Transcript-level abundances were converted to gene-level count estimates using tximport v1.8.0. Differential expression was calculated using DESeq2 v1.20.0. The data were analyzed through IPA (Qiagen). To generate hierarchical clustering, Euclidean distances were calculated using the formula 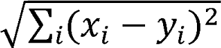 in R.

### Immunoblot validation of RPPA study

Primary UM-HET3 fibroblasts were seeded in six-well plates at 500,000 cells/well 24h prior to drug treatment. Media was replaced by complete DMEM containing 4 µM drug or control (DMSO). After 24 hours drug treatment, cell were lysed in radioimmunoprecipitation assay buffer (RIPA Buffer, Fisher Scientific, Pittsburgh, PA, USA) supplemented with Complete Protease Inhibitor Cocktail (Cat# 04693132001, Sigma Inc. St. Louis, MO) and PhosStop (Cat#4906845001, Sigma Inc. St. Louis, MO). Cell lysates were collected. Protein content was measured using a BCA assay (Cat#23225, Fisher Scientific, Pittsburgh, PA, USA). Protein extracts were fractionated by SDS/PAGE on a 4-15% gel, transferred to PVDF membranes, and electro-transferred to an Immobilon-P Transfer Membrane (Millipore, Billerica, MA, USA) for immunoblot analyses. Membranes were pre-blocked in Tris buffered saline containing 0.05% Tween-20 (TBS-T) and 5% Bovine Serum Albumin (BSA) for 1 hour. After blocking, membranes were probed overnight with primary antibodies in TBS-T supplemented with 5% BSA with shaking at 4°C, followed by three 10 minute washes with TBS-T, incubation with secondary antibody for 1 hour, and three 10 minute washes with TBS-T. Membranes were then developed using an ECL Chemiluminescent Substrate (Cat#32106, Fisher Scientific, Pittsburgh, PA, USA). The following antibodies were used: anti-AMPK (Cat#2603S, 1:1000, Cell Signaling, Danvers, MA), anti-phospho (Thr172)-AMPK (Cat#2535S, 1:1000, Cell Signaling, Danvers, MA), anti-β-actin (Cat#4970S, 1:1000, Cell Signaling, Danvers, MA) and anti-rabbit IgG (Cat#7074, 1:2000, Cell Signaling, Danvers, MA). Quantification was performed using ImageJ software. Data are presented from multiple independent experiments. All data are presented as mean ± SEM. Student’s two tailed *t*-test was used for comparisons of two experimental groups. *P* values were calculated by Student’s *t*-test; *P* □ < □ 0.05 was regarded as significant.

See supplemental methods for additional experimental details, including for the invertebrate studies.

## Supplementary Figure Legends

**Figure S1: Optimization studies of HTS conditions for induction of stress resistance.** (A) MTFs were plated at 10,000 cells/well in a 384-well plate in media consisting of DMEM/0.5% BSA, without serum or antibiotics. All subsequent additions were in DMEM without BSA. Cell viability was assayed using CellTiter-Glo 24 h following addition of 12 mM PQ. The Z’ factor was 0.62. (B) Similar study as in (A) except that the rescue agent curcumin was added 18 hours prior to PQ addition. (C) Partial rescue of MMS-induced cell death by pretreatment with the Nrf2 inducer arsenite. (D) A set of test agents was evaluated using two separate test plates on the same day using PQ as the stress agent. Each symbol represents a separate test compound. Data are represented on a Log10 scale for clarity.

**Figure S2: Representative Kaplan-Meier survival curves of *C. elegans* treated with selected small molecules.**

**Figure S3:** U**n**supervised **hierarchical clustering of RNA-seq samples based on Euclidean distances.** Close relationships between samples are represented in dark blue, and greater distance, in lighter blue. See Methods section for further details.

**Figure S4:** Gene set enrichment analysis (GSEA) for AEG 3482.

**Figure S5. Transcript levels of Nrf2 target genes for 8 compounds evaluated by RNA-seq (related to** Fig. 5C). Only AEG 3482 and cardamonin show induction of Nrf2 transcriptional targets.

**Figure S6: Heatmap depicting 50 proteins changing most in response to compound treatment in RPPA.** Unsupervised hierarchal clustering analysis of the top 50 proteins with the greatest variance from the mean. Red indicates high expression in comparison to the mean. Blue indicates low expression in comparison to the mean. The top 50 proteins with the greatest variance from the mean was calculated by subtracting the mean value of a protein calculated from all samples from the value of a protein in each sample. The top 50 were selected and plotted using pheatmap in R.

**Figure S7. Effects of selected small molecules on HSP expression.** (A) AEG 3482 provokes a weak “Cellular Response to Heat” signature by GSEA. (B) Immunoblot analysis of HSP70 and HSP90 protein expression in MTFs in response to small molecules indicated.

## Supplementary Tables

**Table S1: Table with primary screening data (Cd, MMS, and PQ).** Both raw scores (fluorescence reading/1000) and adjusted values are shown. To calculate the adjusted score, the scores of each batch (day) were multiplied by a scaling factor, so that each batch had the same median score.

**Table S2: Dose response data on 67 compounds successfully tested.** Max, ED50 values for PQ, Cd, and MMS for each compound are shown. P, C, M designations reflect a score of ≥100, at one or more doses, in P, C, and/or M dose response ≥ titrations. Optimally protective doses are shown in the PQ_Max, Cd_Max, and MMS_Max columns. PQy, Cdy, and MMSy columns reflect whether an agent was identified as protective (1) or not protective (0) for that particular stressor.

**Table S3: Lifespan effects of selected small molecules in *C. elegans* and *Drosophila.*** The controls tab in the excel sheet refers to the average mean and median lifespan of the DMSO-fed treatments for that round in the *Drosophila* experiments. A control treatment was performed on every plate; thus if more than 11 drugs were tested during a single round, then more than one control was tested (fly chambers were maintained on 12-well plates), Mean and median were calculated from all of the controls performed in that round.

**Table S4:** Comparison of induction of stress fibroblast resistance and C. elegans **longevity by LOPAC compounds**. Columns L-R are worm data from ^18^.

**Table S5: DrugBank activity annotations for compounds used in primary screening.** Columns in table include: (A) internal small molecule identifier, (B)-(G) activity classification, as in other tables, (H) identifier of nearest structural neighbor found in DrugBank database, (I) Tanimoto coefficient of corresponding structural similarity search, (J) FDA approval status of DrugBank entry, (K) UniProt identifier of corresponding target protein, and (L) pharmacodynamics annotation provided by DrugBank.

**Table S6: Annotations of targets proteins of primary screening compounds.** Table provides more detailed annotation information for target proteins listed in Table S5. Columns include: (A) UniProt identifier, (B)-(C) gene symbol, (D) curation status, (E) functional description of protein, (F) gene name, (G) organism name, (H) length of protein, and (I)-(J) number of FDA and non-FDA approved drugs annotated for corresponding target protein, respectively.

**Table S7: Target set enrichment analysis (TSEA) of target sets identified for active compounds in each stress treatment.** Results for the three functional annotation types GO, KEGG and DO are row-wise appended. Columns include: (A) name of target protein test set; (B) functional annotation type; (C-D) identifier and name of annotation type; (E) number of gene identifiers in the test set annotated to a specific annotation entry divided by the number of gene identifiers in the test set associated with any annotation entry within a given annotation type; (F) same as E but for the universe gene set; (G-I) hypergeometric distribution enrichment test results including raw p-value, adjusted p-value and q-value, respectively; (J) gene identifiers and their (K) quantity matching to corresponding annotation entry in the test set.

**Table S8: Drug set enrichment analysis (DSEA) for KEGG pathways.** The active compound sets from each of the three stress treatments were used for enrichment analysis after mapping them via their target protein annotations to KEGG pathways. The columns in this table are organized the same way as in Table S7 with the exception that the functional annotation type column has been omitted since the analysis included only KEGG pathways.

**Table S9: RNA-seq metrics for 8 compounds tested.** FC, fold change.

**Table S10. IPA immune response-related pathways for the 8 tested compounds.** Pathway analysis of the top 10 immune-related pathways for the 8 compounds. P-values are represented in a -log form and grey values represent non-significant pathways.

**Table S11: Pathways related to oxidative stress response from IPA of DE transcripts.** P-values are represented in a -log format; black font indicates pathways that were significantly induced based on a p-value threshold of 0.05. Grey values = non-significant.

**Table S12: Statistical analysis of RPPA data for 8 compounds for p-AMPK, total AMPK, p-ACC1, and total ACC1.** Median-centered relative intensities were averaged and compared to DMSO values by Student’s T-test. Green indicates significantly increased levels; peach indicates significantly reduced levels.

