## Supplemental figures for "High throughput small molecule screening reveals NRF2-dependent and - independent pathways of cellular stress resistance"

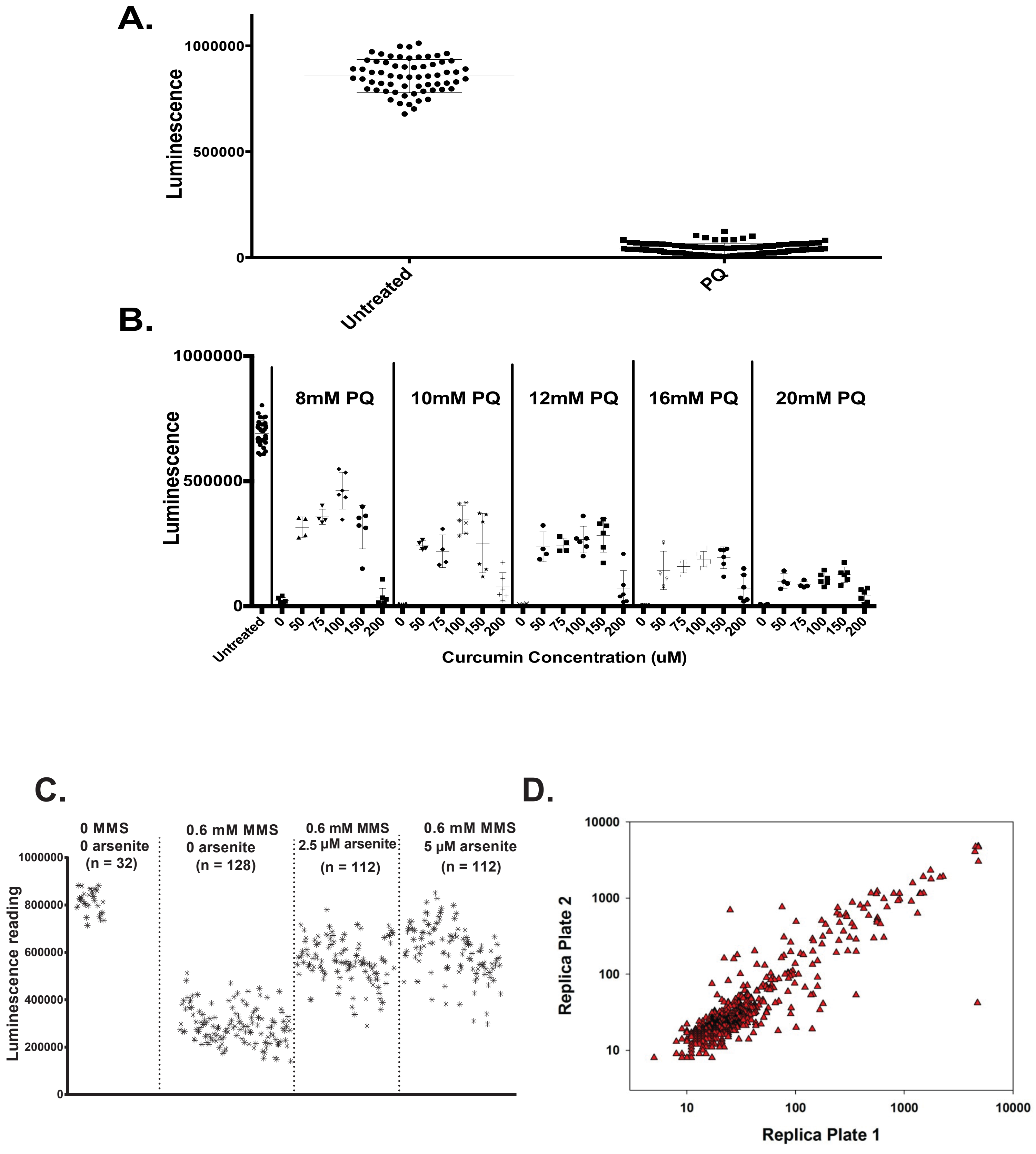

A.
Luminescence
1000000
500000
0
Untreated
PQ
B.
Luminescence
1000000
500000
0
8mM PQ
10mM PQ
12mM PQ
16mM PQ
20mM PQ
Untreated
0
50
75
100
150
200
Curcumin Concentration (uM)
C.
0 MMS
0 arsenite
(n = 32)
0.6 mM MMS
0 arsenite
(n = 128)
0.6 mM MMS
2.5 μM arsenite
(n = 112)
0.6 mM MMS
5 μM arsenite
(n = 112)
Luminescence reading
1000000
800000
600000
400000
200000
0
D.
Replica Plate 2
Replica Plate 1
10
100
1000
10000


Figure S1

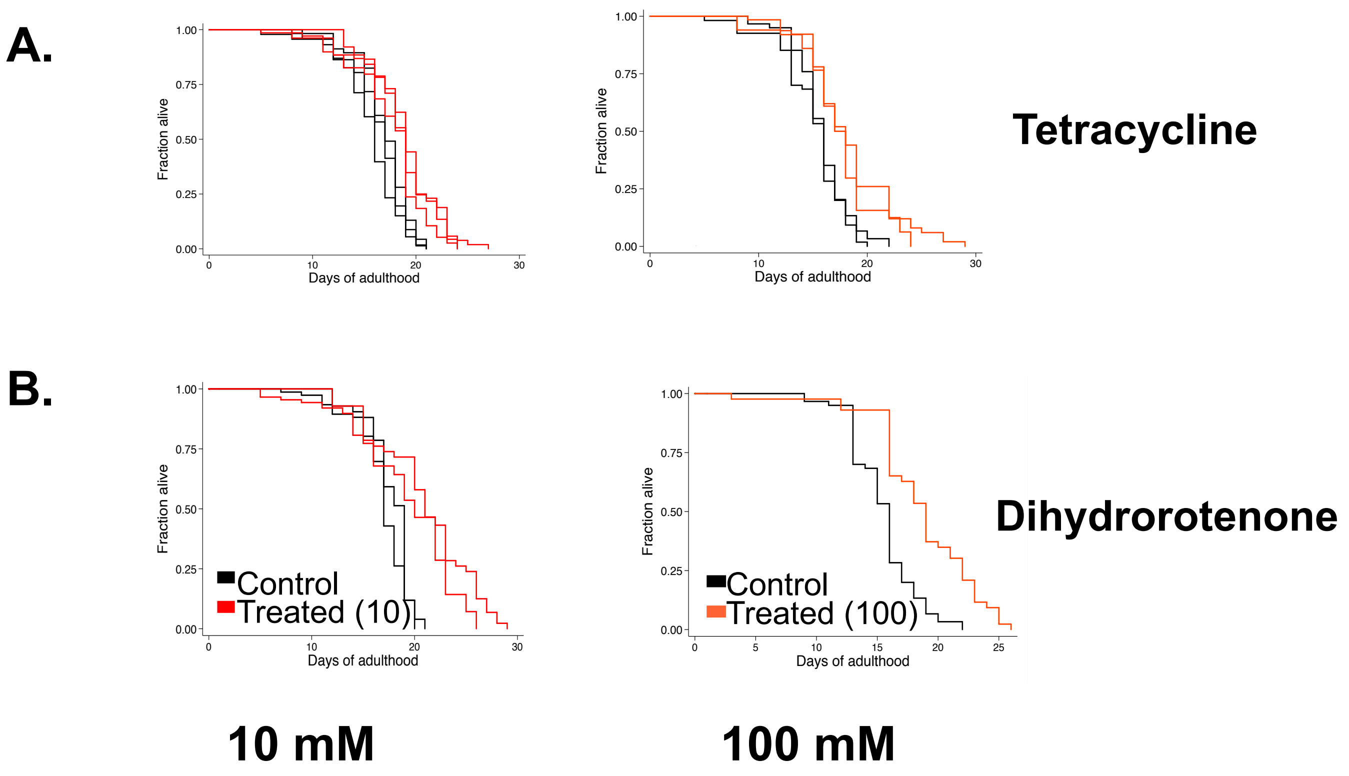
A.
Fraction alive
Days of adulthood
Tetracycline
B.
Control
Treated (10)
Control
Treated (100)
Dihydrorotenone
10 mM
100 mM

### Figure S2

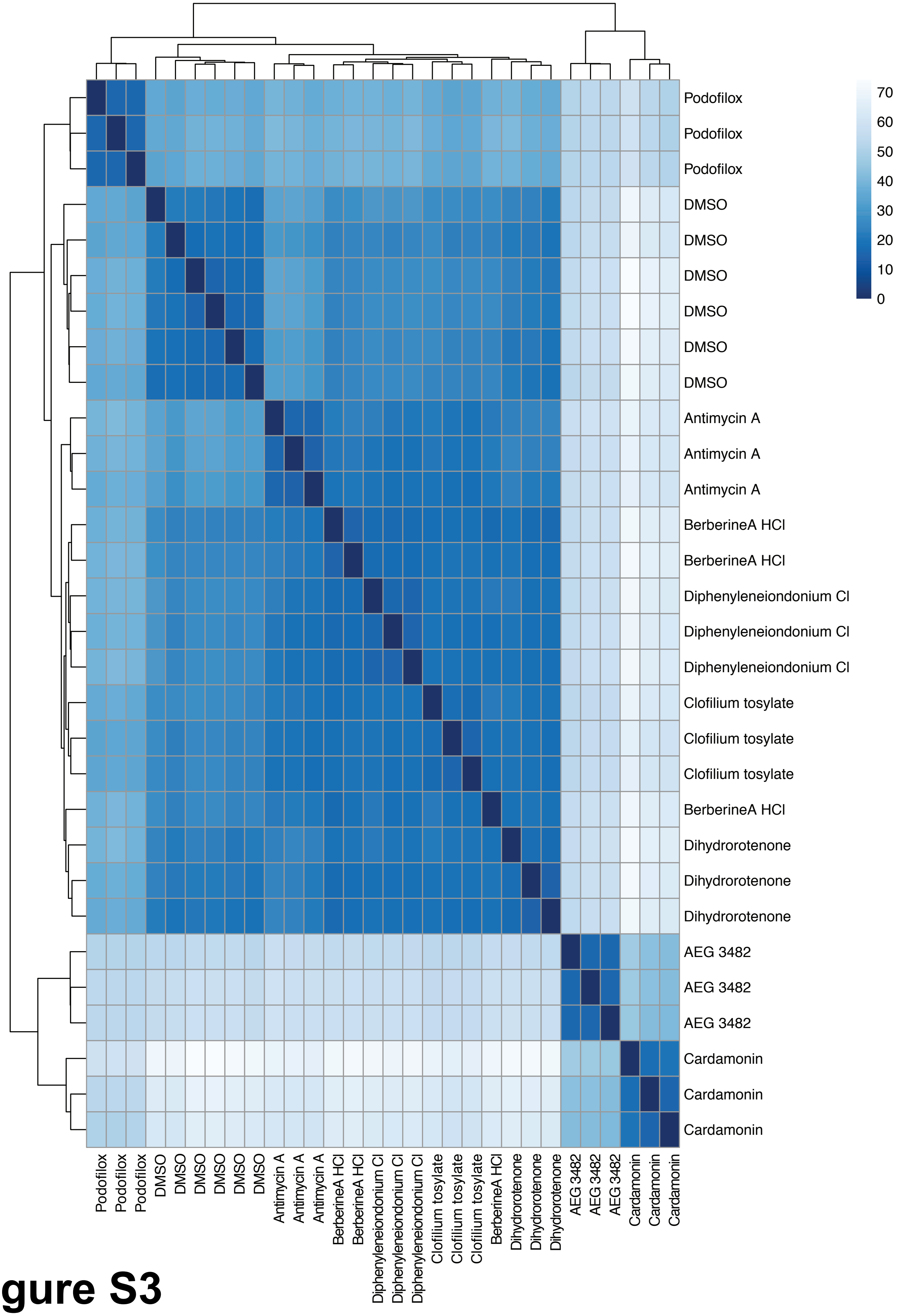

Podofilox
Podofilox
Podofilox
DMSO
DMSO
DMSO
DMSO
DMSO
DMSO
Antimycin A
Antimycin A
Antimycin A
BerberineA HCl
BerberineA HCl
Diphenyleneiondonium Cl
Diphenyleneiondonium Cl
Diphenyleneiondonium Cl
Clofilium tosylate
Clofilium tosylate
Clofilium tosylate
BerberineA HCl
Dihydrorotenone
Dihydrorotenone
Dihydrorotenone
AEG 3482
AEG 3482
AEG 3482
Cardamonin
Cardamonin
Cardamonin
70
60
50
40
30
20
10
0


### Figure S3

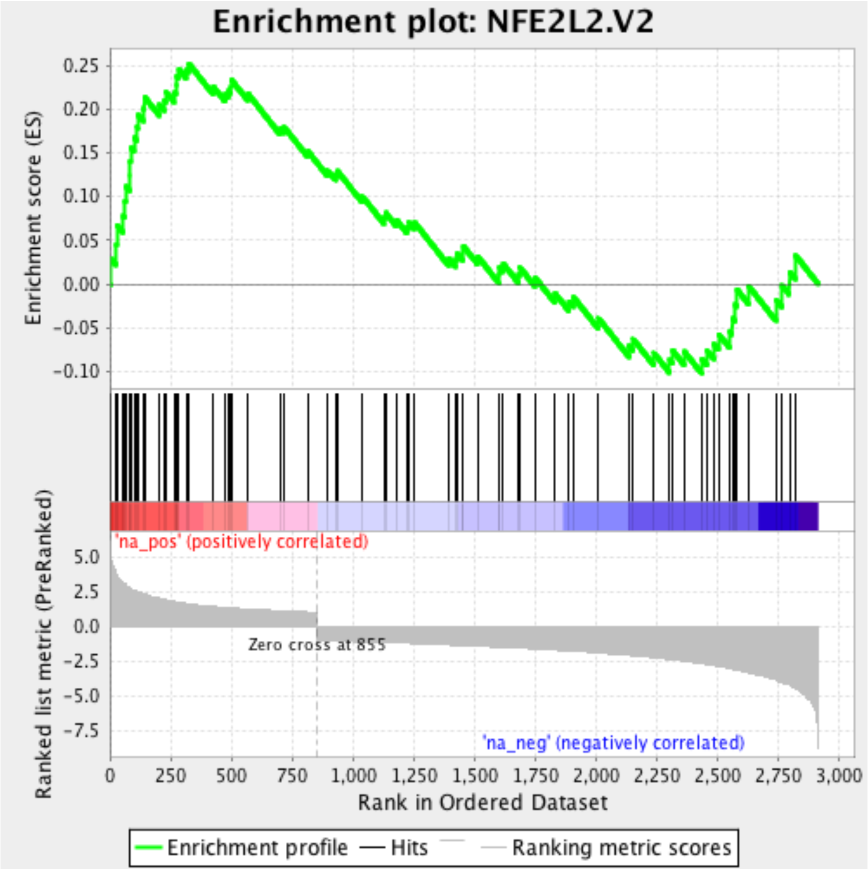


Enrichment score: 0.252
Normalized Enrichment score (NES): 1.90
FDR q-value: 0.148

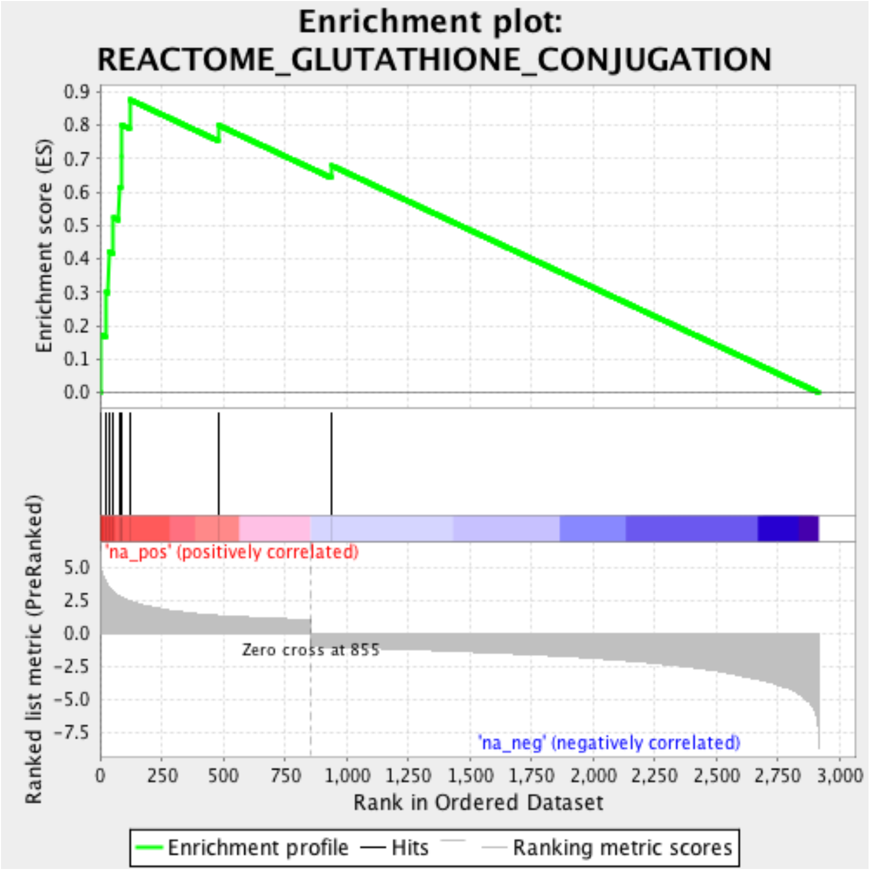


Enrichment score: 0.877
Normalized Enrichment score (NES): 3.11
FDR q-value: 0.00

**Figure S4**

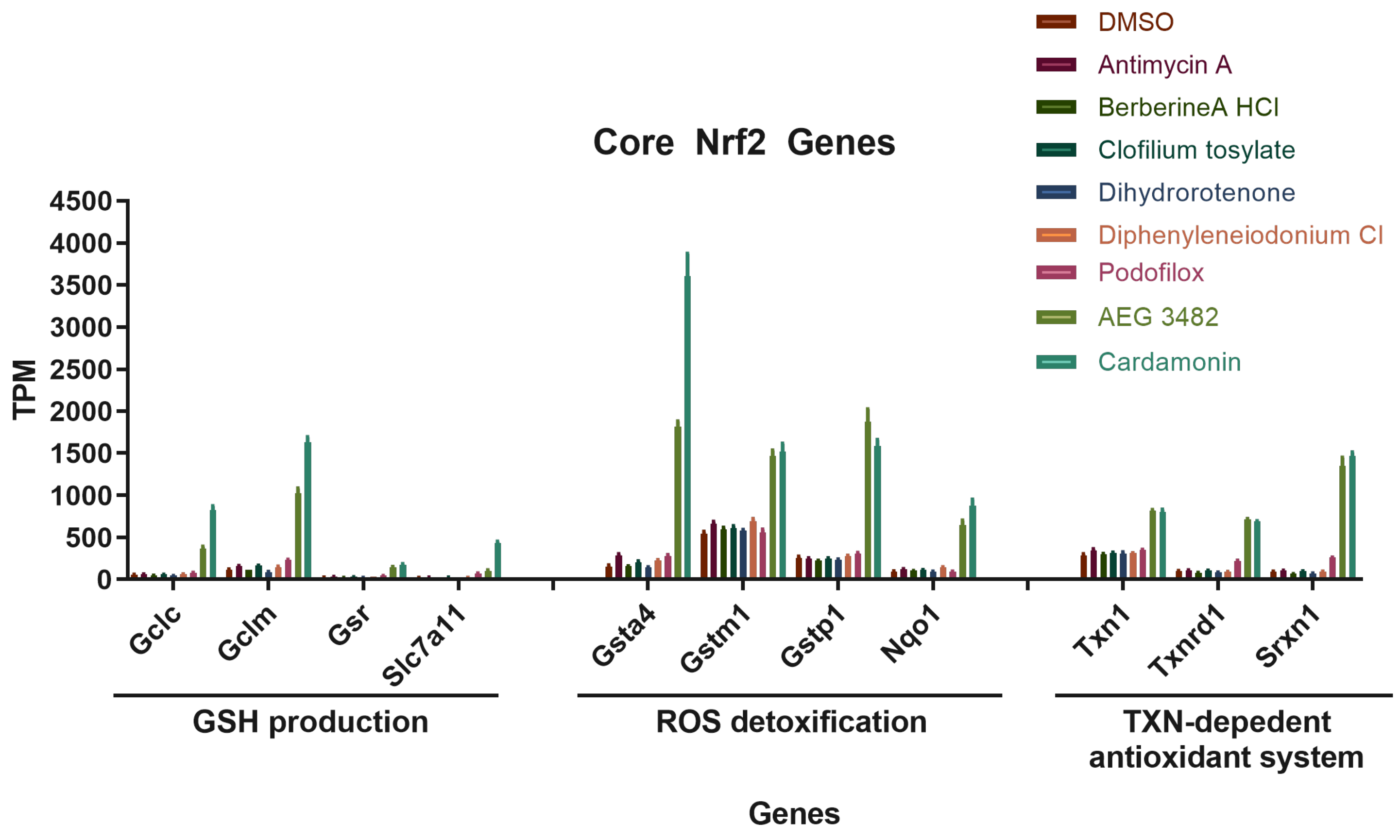
Core Nrf2 Genes
TPM
4500
4000
3500
3000
2500
2000
1500
1000
500
0
Gclc
Gclm
Gsr
Slc7a11
Gsta4
Gstm1
Gstp1
Nqo1
Txn1
Txnrd1
Srxn1
GSH production
ROS detoxification
TXN-depedent antioxidant system
Genes
DMSO
Antimycin A
BerberineA HCl
Clofilium tosylate
Dihydrorotenone
Diphenyleneiodonium Cl
Podofilox
AEG 3482
Cardamonin

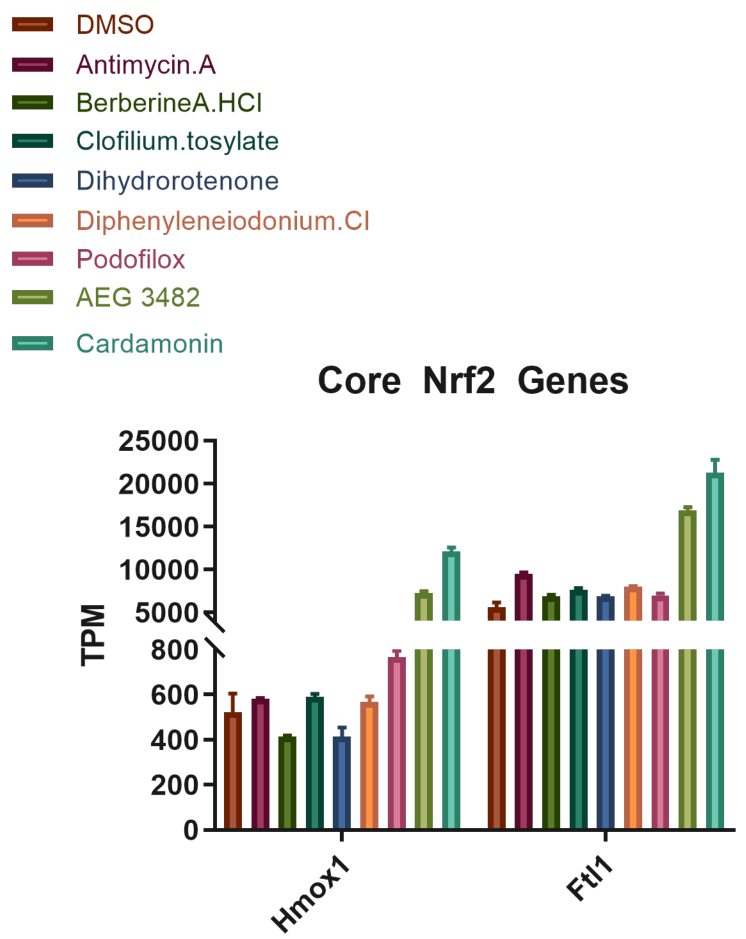
DMSO
Antimycin.A
BerberineA.HCl
Clofilium.tosylate
Dihydrorotenone
Diphenyleneiodonium.Cl
Podofilox
AEG 3482
Cardamonin
Core Nrf2 Genes
TPM
25000
20000
15000
10000
5000
800
600
400
200
0
Hmox1
Ftl1

Figure S5

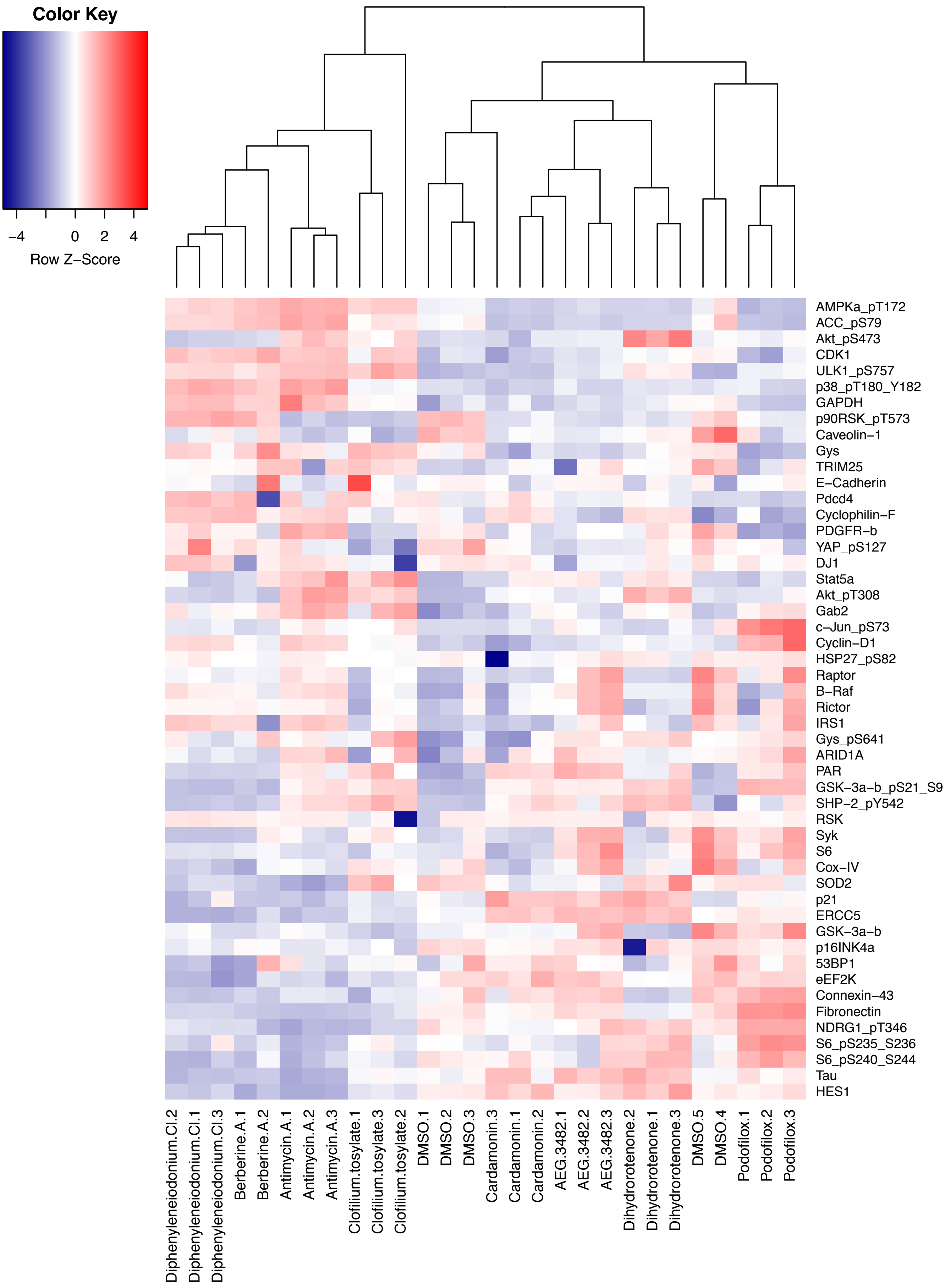

Color Key
−4
0
2
4
Row Z−Score
AMPKa_pT172
ACC_pS79
Akt_pS473
CDK1
ULK1_pS757
p38_pT180_Y182
GAPDH
p90RSK_pT573
Caveolin−1
Gys
TRIM25
E−Cadherin
Pdcd4
Cyclophilin−F
PDGFR−b
YAP_pS127
DJ1
Stat5a
Akt_pT308
Gab2
c−Jun_pS73
Cyclin−D1
HSP27_pS82
Raptor
B−Raf
Rictor
IRS1
Gys_pS641
ARID1A
PAR
GSK−3a−b_pS21_S9
SHP−2_pY542
RSK
Syk
S6
Cox−IV
SOD2
p21
ERCC5
GSK−3a−b
p16INK4a
53BP1
eEF2K
Connexin−43
Fibronectin
NDRG1_pT346
S6_pS235_S236
S6_pS240_S244
Tau
HES1
Diphenyleneiodonium.Cl.2
Diphenyleneiodonium.Cl.1
Diphenyleneiodonium.Cl.3
Berberine.A.1
Berberine.A.2
Antimycin.A.1
Antimycin.A.2
Antimycin.A.3
Clofilium.tosylate.1
Clofilium.tosylate.3
Clofilium.tosylate.2
DMSO.1
DMSO.2
DMSO.3
Cardamonin.3
Cardamonin.1
Cardamonin.2
AEG.3482.1
AEG.3482.2
AEG.3482.3
Dihydrorotenone.2
Dihydrorotenone.1
Dihydrorotenone.3
DMSO.5
DMSO.4
Podofilox.1
Podofilox.2
Podofilox.3


### Figure S6

A.

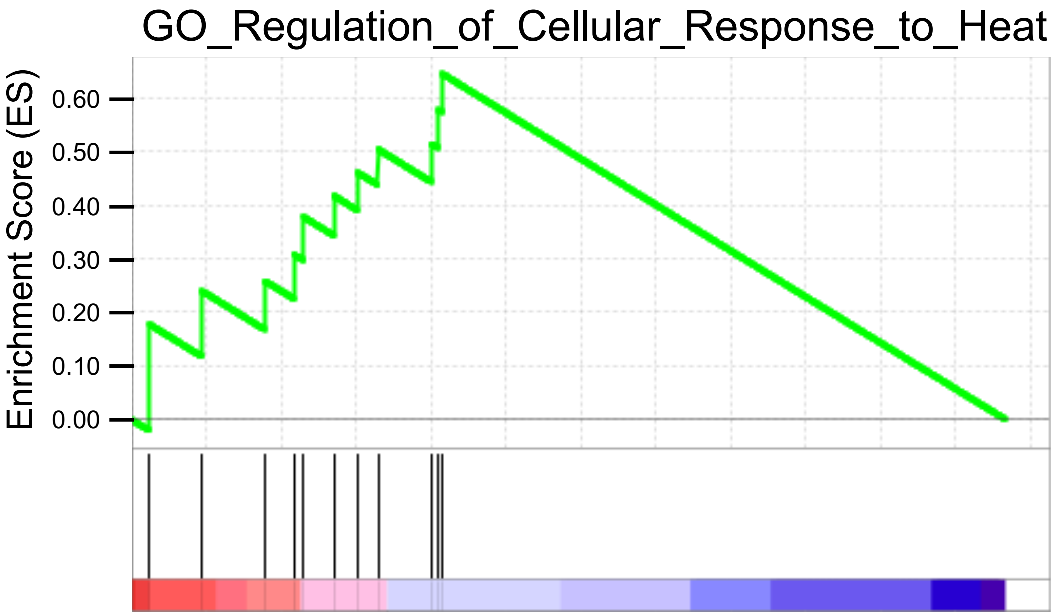
GO_Regulation_of_Cellular_Response_to_Heat
Enrichment Score (ES)
0.60
0.50
0.40
0.30
0.20
0.10
0.00

B.

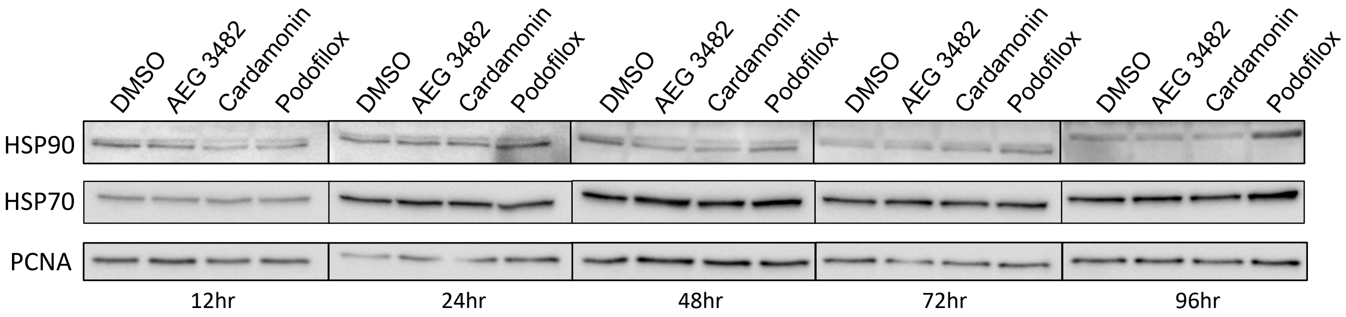
DMSO
AEG 3482
Cardamonin
Podofilox
HSP90
HSP70
PCNA
12hr
24hr
48hr
72hr
96hr

### Figure S7
