## Supplemental methods for "High throughput small molecule screening reveals NRF2-dependent and - independent pathways of cellular stress resistance"

**Supplementary Methods**

**Drosophila *lifespan measurements:***

A wild type laboratory strain of *D. melanogaster*, Canton S, was obtained from the Bloomington Stock Center for all experiments. To prepare flies with similar developmental environments, larvae were cultured under same density conditions in cornmeal-sugar-yeast media in standard light/dark conditions at 25°C. Same-age adults were then collected within 24 hours of emergence and transferred onto standard sugar-yeast (SY) food for 48-72 hours to allow for mating. After this time, 20 flies were sorted by sex into vials under light CO_2_ anesthetic. The vials used in these experiments were created within the laboratory to allow for high-throughput analysis and consisted of 12 standard fly vials glued together to create a rectangle (3 vials high by 4 vials wide). Vials were screened on one end to allow for air exchange and open on the other end to allow the flies free access to the food. This design enabled the use of standard 12-well culture plates (Corning Costar) as the reservoir of drug/food that could easily be exchanged by simply tapping the flies down onto the screen side of the vials, removing the old 12-well plate, and replacing with a new 12-well plate that contains fresh drug/food.

All of the chemicals used in the lifespan experiments were obtained either from Sigma-Aldrich or Fisher Scientific. The chemicals were prepared at 10 mM concentrations in DMSO (stock solutions) and stored at -20°C in several aliquots, so that no chemical was frozen/thawed more than 3 times. The stock solutions were thawed and diluted into water at 20 μM for females or 200 μM for males, since male flies eat less food than female flies. A control vial, containing just DMSO diluted in water, was included in every 12-well group. The chemical concentrations were chosen based upon empirical work in the lab as well as upon the concentrations of other drugs reported to extend lifespan in *Drosophila*. The diluted solutions were then coated on top of standard SY food and allowed to dry. All chemical handling was done using an Eppendorf epMotion 5075 to automate the process. The placement of each chemical/control was randomized across the 12-well plate prior to the start of the experiment and then remained constant for the entire experiment thereafter. We administered each chemical to 4 separate vials, across 4 different 12-well plates, resulting in a total of 80 flies/treatment/sex. The freshly prepared food was given to the flies every Monday, Wednesday, and Friday. At the time, the number of dead flies were counted and recorded using Dlife software^1^. All of the survivorship data were analyzed using a Log-Rank test between the chemical of interest and the DMSO control.

**C. elegans *lifespan measurements:***

**Plate preparation:** The screenings for longevity were carried out in 12-well plate with 3.5 mL of NG agar added to each well. The plates were allowed to solidify and dried for at least 48 hours before 10 μL of *E. Coli* OP50 was added into each well. The plates were then transferred to 37°C overnight. Finally, the OP50 bacteria were killed by 30 min of UV light, before 3.5 μL of drug stock solution (dissolved in DMSO unless indicated otherwise) were added into each well. The concentrations indicated in the paper is the final concentration for each well (final volume 3.5 mL). Compounds were added to the NG plates at least 24 hours before transferring animals onto drug-containing plates. The final DMSO concentration was kept at 0.1% after adding drugs to the plates.

**Worm preparation:** The screening was carried out using the sterile strain, CF512: *fer-15(b26)II; fem-1(hc17)III*^2^. All worms used were maintained at 15°C and handled as described previously^3^. Eggs of CF512 worms were harvested from young adults and seeded in regular NG plates to be synchronized. Synchronized Day 1 adults were collected from plates and washed in M9 5 times before transferred onto the drug-containing screening plates mentioned above. Approximately 60 worms were placed in each well and grown at 25°C. To ensure the drugs retain potency throughout the entire experiment and to avoid starvation, bacteria food and drug were replenished every 5 days. Viability of the worms were scored every 1-3 days until 100% of the animals died.

**Lifespan statistics:** In all experiments, the pre-fertile period of adulthood was used as t = 0 for lifespan analysis. RStudio software was used for statistical analysis to determine the means and P-value. In all cases, P values were calculated using the stratified log-rank test.

| **CCG #** | **Compound** | **CAS #** | **Vendor** | **Catalog #** | **Smiles Structure** |
| --- | --- | --- | --- | --- | --- |
| CCG-39894 | PODOFILOX (Podophyllotoxin in Sigma) | 518-28-5 | Sigma | P4405 | COc1cc(cc(OC)c1OC)[C@H]1[C@@H]2[C@H](COC2=O)[C@@H](O)c2cc3OCOc3cc12 |
| CCG-220222 | Antimycin A | 1397-94-0 | Sigma | A8674 | CCCCCC[C@@H]1[C@@H](OC(=O)CC(C)C)[C@H](C)OC(=O)C(NC(=O)c2cccc(NC=O)c2O)[C@@H](C)OC1=O |
| CCG-221541 | Supercinnamaldehyde | 703-51-8 | Sigma | S3322 | CN1C(=O)\C(=C/C(C)=O)c2ccccc12 |
| CCG-38389 | DIHYDROROTENONE | ** | Aldrich CPR | ** | [H][C@@]12COc3cc(OC)c(OC)cc3[C@]1([H])C(=O)c1ccc3OC(Cc3c1O2)C(C)C |
| CCG-208127 | Cardamonin | 19309-14-9 | Fisher | 250910 | COc1cc(O)cc(O)c1C(=O)\C=C\c1ccccc1 |
| CCG-208552 | BerberineA HCl | 633-65-8 | Sigma | B3251-5G | Cl.COc1ccc2cc3-c4cc5OCOc5cc4CC[n+]3cc2c1OC |
| CCG-220319 | Clofilium tosylate | 92953-10-1 | Fisher | ICN15374925 | Cc1ccc(cc1)S([O-])(=O)=O.CCCCCCC[N+](CC)(CC)CCCCc1ccc(Cl)cc1 |
| CCG-221955 | AEG 3482 | 63735-71-7 | Fisher | 26-511-0 | NS(=O)(=O)c1nn2cc(nc2s1)-c1ccccc1 |

**Compounds used in RNA-seq and RPPA analysis:** The sources and catalog numbers for the compounds used in RNA-seq and RPPA are as follows:

** Dihydrorotenone was obtained as a special order through AldrichCPR.

**Supplemental Methods references:**

1.	Linford, N.J., Bilgir, C., Ro, J. & Pletcher, S.D. Measurement of lifespan in Drosophila melanogaster. *J Vis Exp* (2013).

2.	Garigan, D. et al. Genetic analysis of tissue aging in Caenorhabditis elegans: a role for heat-shock factor and bacterial proliferation. *Genetics* **161**, 1101-12 (2002).

3.	Brenner, S. The genetics of Caenorhabditis elegans. *Genetics* **77**, 71-94 (1974).
